# Mitochondrial DNA Variations Modulate Alveolar Epithelial Mitochondrial Function and Oxidative Stress in Newborn Mice Exposed to Hyperoxia

**DOI:** 10.1101/2023.05.17.541177

**Authors:** Jegen Kandasamy, Rui Li, Bianca M. Vamesu, Nelida Olave, Brian Halloran, Tamas Jilling, Scott W Ballinger, Namasivayam Ambalavanan

**Affiliations:** Department of Pediatrics, University of Alabama at Birmingham School of Medicine, Birmingham, Alabama; Department of Pediatrics, University of South Alabama, Mobile, Alabama; Department of Pathology, University of Alabama at Birmingham School of Medicine

**Author notes:** **Corresponding Author:** Jegen Kandasamy, Department of Pediatrics, University of Alabama at Birmingham School of Medicine, 1700 6th Avenue South, Birmingham, AL 35233.

## Abstract

Oxidative stress is an important contributor to bronchopulmonary dysplasia (BPD), a form of chronic lung disease that is the most common morbidity in very preterm infants. Mitochondrial functional differences due to inherited and acquired mutations influence the pathogenesis of disorders in which oxidative stress plays a critical role. We previously showed using mitochondrial-nuclear exchange (MNX) mice that mitochondrial DNA (mtDNA) variations modulate hyperoxia-induced lung injury severity in a model of BPD. In this study, we studied the effects of mtDNA variations on mitochondrial function including mitophagy in alveolar epithelial cells (AT2) from MNX mice. We also investigated oxidant and inflammatory stress as well as transcriptomic profiles in lung tissue in mice and expression of proteins such as PINK1, Parkin and SIRT3 in infants with BPD. Our results indicate that AT2 from mice with C57 mtDNA had decreased mitochondrial bioenergetic function and inner membrane potential, increased mitochondrial membrane permeability and were exposed to higher levels of oxidant stress during hyperoxia compared to AT2 from mice with C3H mtDNA. Lungs from hyperoxia-exposed mice with C57 mtDNA also had higher levels of pro-inflammatory cytokines compared to lungs from mice with C3H mtDNA. We also noted changes in KEGG pathways related to inflammation, PPAR and glutamatergic signaling, and mitophagy in mice with certain mito-nuclear combinations but not others. Mitophagy was decreased by hyperoxia in all mice strains, but to a greater degree in AT2 and neonatal mice lung fibroblasts from hyperoxia-exposed mice with C57 mtDNA compared to C3H mtDNA. Finally, mtDNA haplogroups vary with ethnicity, and Black infants with BPD had lower levels of PINK1, Parkin and SIRT3 expression in HUVEC at birth and tracheal aspirates at 28 days of life when compared to White infants with BPD. These results indicate that predisposition to neonatal lung injury may be modulated by variations in mtDNA and mito-nuclear interactions need to be investigated to discover novel pathogenic mechanisms for BPD.

## 1. Introduction

Oxidative stress is a major risk factor for bronchopulmonary dysplasia (BPD) which is a significant cause of morbidity and mortality in very premature extremely low birth weight (ELBW) infants. BPD-associated accelerated cardiopulmonary aging also frequently leads to persistent lung dysfunction during adult life [1,2]. A growing body of evidence indicates mitochondrial function plays a critical role in oxidative stress pathophysiology that contribute to lung disorders such as chronic obstructive pulmonary disease (COPD) and idiopathic pulmonary fibrosis (IPF) [3]. Similarly, inherited mitochondrial DNA (mtDNA) polymorphisms (haplogroups) modulate risk for lung diseases such as pulmonary hypertension (PH) and asthma in adults [4,5]. Mitochondrial function spans several critical aspects of cellular homeostasis during oxidative stress. ATP generation through electron transport chain (ETC) function and oxidative phosphorylation provides energy required for maintenance of overall cellular heath [6]. Mitochondria are also a major source of oxidants that damage vital intracellular structures, including mitochondrial components such as mtDNA. Mitochondrial antioxidant enzymes such as aconitase and SIRT3 are essential in combating such oxidant stress [7,8]. When quality control mechanisms such as mitophagy are unable to re-establish mitochondrial functional integrity, increased mitochondrial permeability transition pore (MPTP) opening and decreased mitochondrial membrane potential (ΔΨm) can initiate cell death mechanisms such as apoptosis [9,10]. Such changes in type II alveolar epithelial cells (AT2) and lung fibroblast mitochondrial function and quality control are known to increase lung inflammation and injury in mice and in individuals with COPD, IPF and asthma [11–13].

We have previously found that human umbilical venous endothelial cell (HUVEC) and mesenchymal stem cell (MSC) mitochondrial function predicts risk for BPD in ELBW infants [14,15]. We have also found that mice carrying C57 mtDNA have increased lung injury and neonatal murine lung fibroblast (NMLF) mitochondrial dysfunction when compared to mice carrying C3H mtDNA in a hyperoxia-induced BPD model using wild-type C57 (C57WT, with C57^n (nuclear)^ C57^mt(mitochondrial)^ genotype; more sensitive to hyperoxia-induced lung injury) and C3HWT (C3H^n^C3H^mt^; more resistant to hyperoxic lung injury) mice as well as their corresponding mitochondrial-nuclear exchange (MNX) mice strains C57MNX (C57^n^C3H^mt^) and C3HMNX (C3H^n^C57^mt^) [16]. However, we did not investigate whether such differences in mitochondrial function that modulate lung injury severity in ELBW infants with BPD as well as in mice with different mtDNA are also associated with increased mitochondrial oxidant stress and deterioration of quality control mechanisms such as mitophagy. We also have not previously studied the impact of mtDNA variations on hyperoxia-induced mitochondrial functional changes in cell types such as AT2 which play key roles in lung development.

Therefore, in this study, we examined the effects of differences in mtDNA haplogroups on mitochondrial function and oxidative stress in AT2 from newborn WT and MNX mice exposed to normoxia or hyperoxia. Next, we measured markers of oxidative stress and inflammation and used RNA sequencing to study changes in the lung transcriptome in the lungs of these mice. Finally, we also evaluated quality control mechanisms such as mitophagy in the lungs, AT2 and NMLF of WT and MNX mice exposed to normoxia or hyperoxia, as well as in HUVEC and tracheal aspirates (TA) obtained from a cohort of White and Black ELBW infants with moderate/severe BPD.

## 2. Results

### 2.1. MtDNA modulates hyperoxia-induced AT2 bioenergetic dysfunction

Bioenergetic function of intact and permeabilized AT2 cells isolated from WT and MNX mice exposed to normoxia or hyperoxia was determined by measuring oxygen consumption rate (OCR) using an Oroboros Oxygraph. Hyperoxia exposure decreased AT2 basal OCR in C57WT, C57MNX and C3HMNX mice (mean [SD], pmol/sec/10^6^ cells, 63 [8] vs. 123 [22], 94 [10] vs. 131 [16], and 74 [6] vs. 125 [13] respectively, p <0.005 for all comparisons) but not in C3HWT mice (101 [9] vs. 124 [25], p = 0.09). Basal OCR of AT2 cells exposed to normoxia was similar between all groups. When exposed to hyperoxia, basal OCR was lower in mice with C57 mtDNA (C57WT and C3HMNX) compared to their nuclear counterparts that carry C3H mtDNA (C57MNX and C3HWT respectively), as shown in **Figure 1A**. Hyperoxia also decreased AT2 maximal OCR (measured in the presence of the protonophore FCCP) in all mice strains (mean [SD], pmol/sec/10^6^ cells, 149 [10] vs. 190 [25] in C57WT, 160 [20] vs. 218 [25] in C57MNX, 172 [16] vs. 212 [23] in C3HWT and, 154 [5] vs. 199 [12] in C3HMNX, p <0.005 for all comparisons). AT2 maximal OCR was similar between normoxia-exposed C57WT vs. C57MNX mice and between C3HWT vs. C3HMNX mice. When exposed to hyperoxia, AT2 maximal OCR continued to remain similar between C57WT vs. C57MNX mice but was lower in C3HMNX mice compared to C3HWT mice (**Figure 1B**).

**Figure 1:**
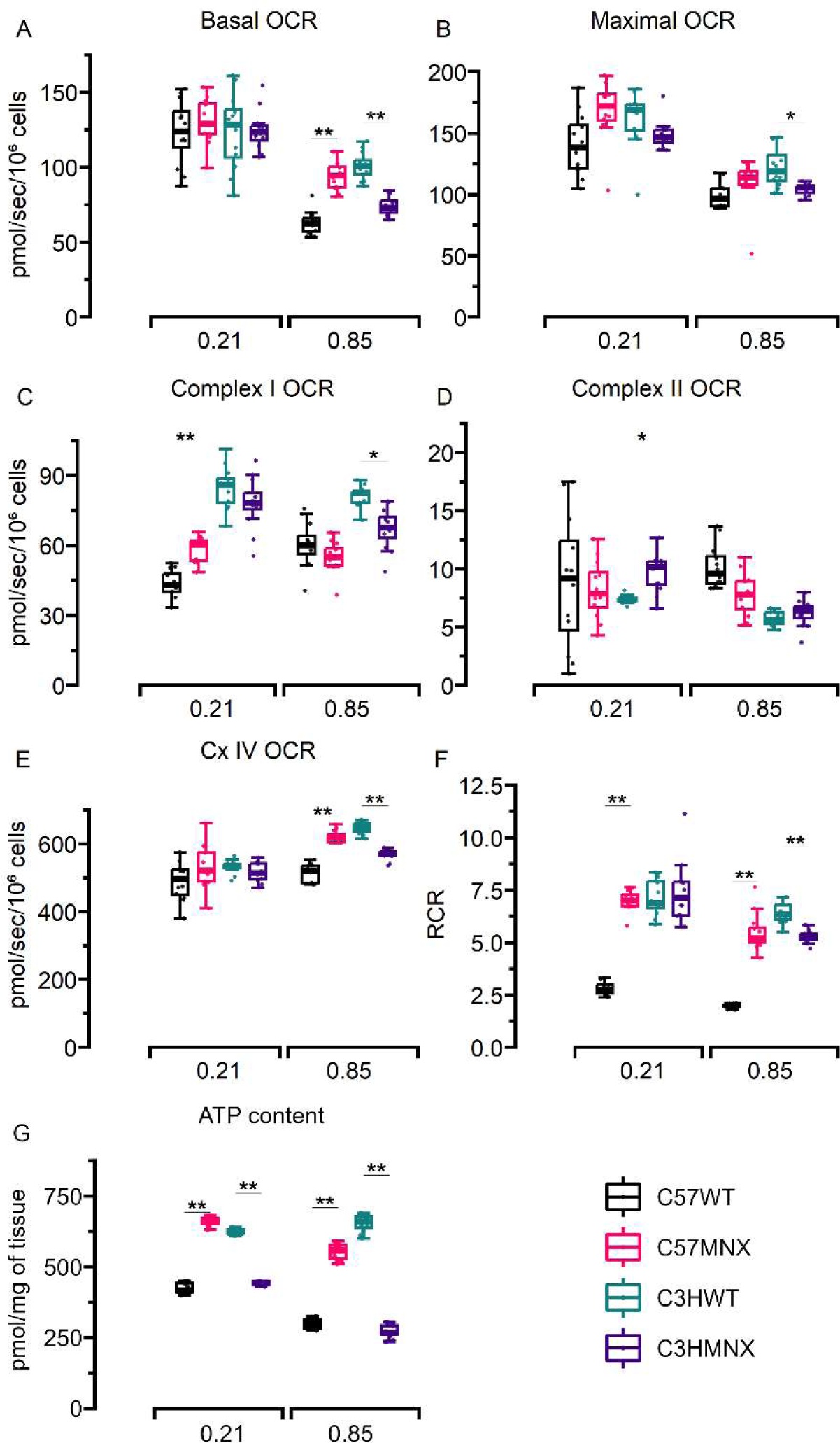
Bioenergetics of intact and permeabilized AT2 cells from mice. **A**: Basal OCR. **B**: Maximal OCR. **C**: Complex I OCR. **D**: Complex II OCR. **E**: Complex IV OCR. **F**: RCR. **G**: ATP content. N = minimum of 12 mice/group. All data were analyzed by 2-way ANOVA or Kruskal-Wallis tests, followed by post hoc analyses. Box represents median/interquartile range, whiskers represent maximum and minimum values. * and ** represent p-values < 0.05 and < 0.005 respectively.

Hyperoxia also caused changes in individual ETC bioenergetic function, most notably for complex IV OCR, which was lower in permeabilized AT2 obtained from hyperoxia-exposed C57WT compared to C57MNX mice and in AT2 from C3HMNX vs. C3HWT mice. Similar changes were also noted in respiratory control ratio (RCR), which is considered the single most useful measure of mitochondrial function [17]. Additionally (and consistent with the generalized hyperoxia-induced decrease in AT2 OCR) ATP content in AT2 lysates was also decreased by hyperoxia in all mice strains (mean [SD], pmol/mg of tissue, 297 [16] vs. 423 [18] in C57WT, 554 [27] vs. 661 [14] in C57MNX, 654 [30] vs. 623 [9] in C3HWT and, 272 [21] vs. 442 [6] in C3HMNX, p<0.05 for all comparisons). However, (and unlike basal OCR which remained similar between groups) ATP content in AT2 was lower in C57WT compared to C57MNX and in C3HMNX compared to C3HWT mice during normoxia. Finally, ATP content was also noted to be lower in AT2 from hyperoxia-exposed mice with C57 mtDNA compared to mice with C3H mtDNA (**Figures 1C-G**). These results suggest that while mitochondrial bioenergetics may not be modified by mtDNA variations during normoxia, hyperoxia-induced mitochondrial bioenergetic dysfunction in alveolar epithelial cells from newborn mice may vary with differences in mtDNA.

### 2.2. AT2 from hyperoxia-exposed mice with C57 mtDNA exhibit higher MPTP permeability and lower ΔΨm

Hyperoxia increased MPTP opening (which was measured by calculating ionomycin-induced calcein signal quenching rate) in AT2 cells from all strains (mean [SD], calcein quenching rate relative to normoxia-exposed C57WT group, 2 [0.3] vs. 1 [0.1] in C57WT, 0.6 [0.1] vs. 0.1 [0.01] in C57MNX, 0.2 [0.02] vs. 0.3 [0.1] in C3HWT and, 1.2 [0.2] vs. 0.28 [0.06] in C3HMNX mice, p < 0.001 for all groups). Under normoxic conditions, MPTP opening was higher in AT2 from C57WT mice compared to C57MNX mice, but similar in C3HMNX vs. C3HWT mice. When exposed to hyperoxia, MPTP opening continued to be higher in AT2 from C57WT mice compared to C57MNX mice and increased to be higher in AT2 from C3HMNX mice compared to C3HWT mice (**Figures 2A, C-D**).

**Figure 2:**
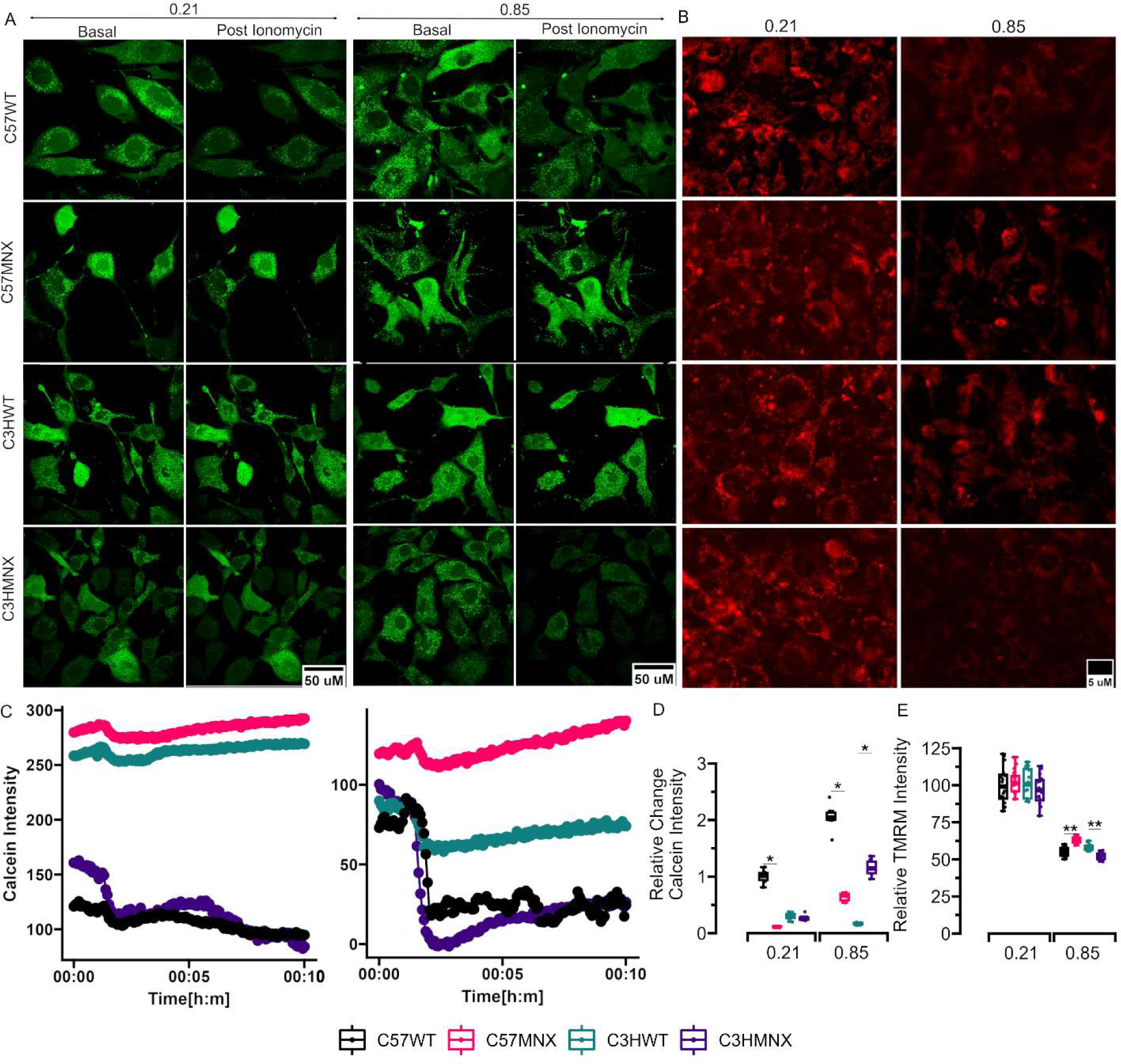
MPTP permeability and ΔΨm measured using calcein-CO signal quenching and TMRM fluorescence in AT2 from mice. **A**: Representative images of AT2 from mice that were stained with calcein and exposed to normoxia/hyperoxia and saline/ionomycin. **B**: Representative images of AT2 from mice that were stained with TMRM and exposed to normoxia/hyperoxia. **C**: Calcein intensity in AT2 measured over time depicting fall in intensity noted after ionomycin was added at 1 minute. **D**: Calcein intensity and **E**: TMRM fluorescence. N = minimum of 6 mice/group. All data were analyzed by 2-way ANOVA or Kruskal-Wallis tests, followed by post hoc analyses. Box represents median/interquartile range, whiskers represent maximum and minimum values. * and ** represent p-values < 0.05 and < 0.005 respectively.

We quantified ΔΨm and depolarization of the inner mitochondrial membrane in AT2 by measuring differences in TMRM fluorescence intensity. Hyperoxia exposure decreased ΔΨm in all mice strains (mean [SD], TMRM fluorescence relative to normoxia-exposed C57WT group, 55 [3] vs. 100 [13] in C57WT, 63 [2] vs. 103 [8] in C57MNX, 58 [2] vs. 101 [10] in C3HWT and, 52 [2] vs. 97 [11] in C3HMNX mice, P < 0.001 for all comparisons). Though TMRM fluorescence was similar between AT2 from normoxia-exposed C57WT vs. C57MNX and C3WHT vs. C3HMNX mice, it was noted to be lower in AT2 from hyperoxia-exposed C57WT mice when compared to C57MNX mice and in AT2 from hyperoxia-exposed C3HMNX vs. C3HWT mice (**Figures 2B, 2E**). Increased inner mitochondrial membrane permeability is associated with both apoptotic and necrotic cell death [18]. In this context, it was notable that the exacerbation of hyperoxia-induced mitochondrial bioenergetic dysfunction noted in AT2 from mice with C57 mtDNA was also accompanied by increased permeabilization as well as depolarization of the mitochondrial membrane.

### 2.3. Hyperoxia-induced AT2 oxidative damage is modified by mtDNA variation

MtDNA is known to be exquisitely sensitive to oxidative stress [19]. To determine whether the AT2 mitochondrial dysfunction that was noted in hyperoxia-exposed mice was associated with increased oxidative stress we measured mtDNA damage in AT2 and found that hyperoxia exposure increased AT2 mtDNA damage in C57WT, C57MNX and C3HMNX mice (mean [SD], lesions/10 kb, 1 [0.2] vs 0.3 [0.1], 0.4 [0.2] vs 0.06 [0.02] and, 0.9 [0.3] vs 0.3 [0.1] respectively, P < 0.005 for all comparisons) but not in C3HWT mice (0.5 [0.2] vs 0.3 [0.1], P = 0.12). When exposed to normoxia, AT2 mtDNA damage was higher in C57WT mice compared to C57MNX mice but similar in C3HMNX vs. C3HWT mice. However, during hyperoxia exposure, AT2 mtDNA damage was higher in mice carrying C57 mtDNA (C57WT and C3HMNX) than their nuclear counterparts (C57MNX and C3HWT respectively) with C3H mtDNA (**Figure 3A**).

**Figure 3:**
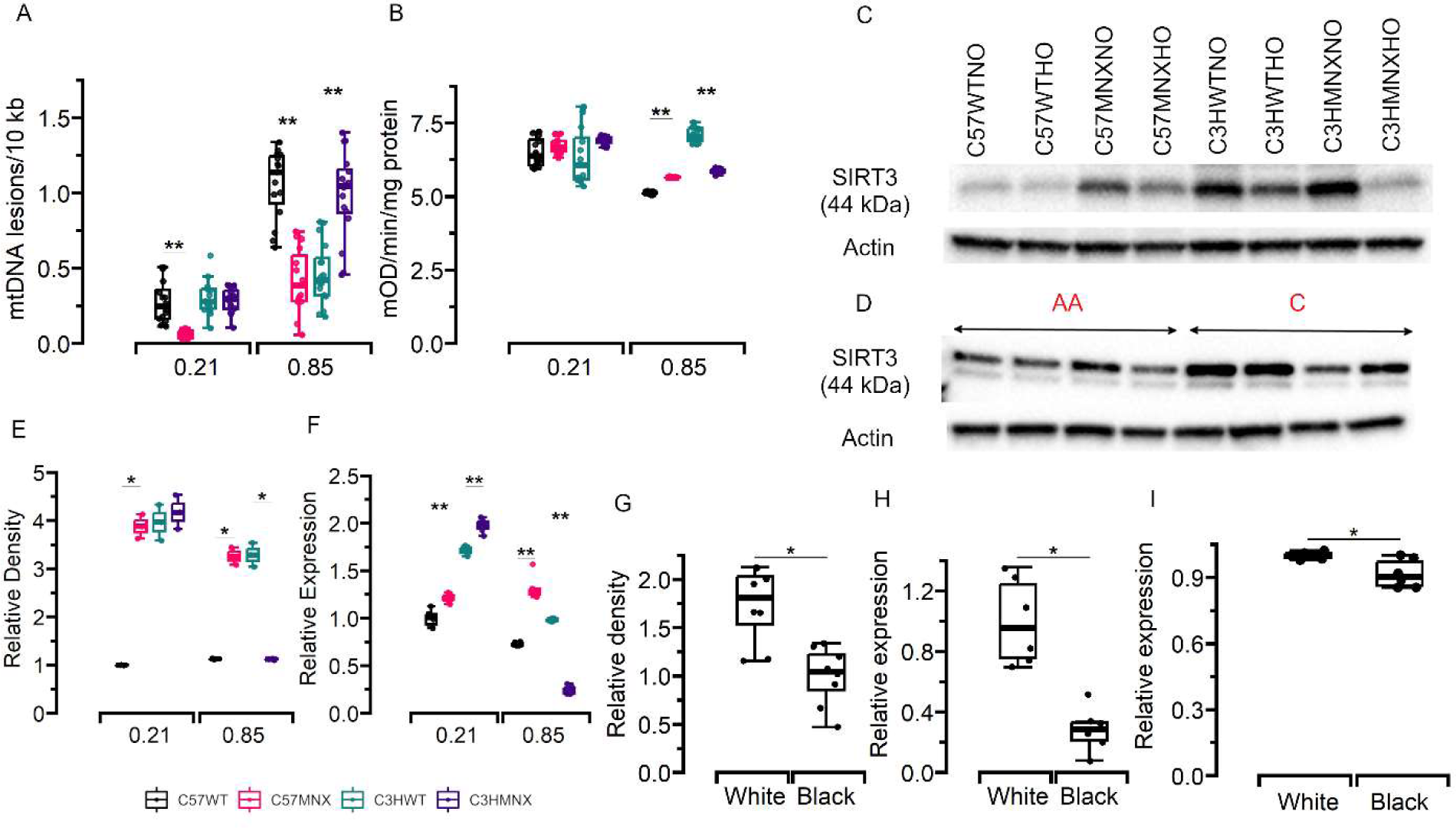
MtDNA damage, aconitase activity and SIRT3 expression in AT2 cells from mice and HUVEC and TA from infants. **A**: MtDNA damage in mice AT2 cells. **B**: AT2 aconitase activity. **C**: Representative Western blots for SIRT3 obtained from AT2 cells. **D**: Representative Western blot for SIRT3 from HUVEC. **E-F**: Densitometry plot and qPCR for SIRT3 from AT2 cells. **G-H**: SIRT3 protein and mRNA expression in HUVEC from White and Black infants who later developed BPD. **I**: PINK1 qPCR in tracheal aspirates obtained around 28 days of life from infants who later developed BPD. N = 6 mice/group and 8 infants per group. All data were analyzed by 2-way ANOVA or Kruskal-Wallis tests, followed by post hoc analyses. Box represents median/interquartile range, whiskers represent maximum and minimum values. * and ** represent p-values < 0.05 and < 0.005 respectively.

Mitochondrial aconitase activity is also considered to be a sensitive and critical target of oxidative stress in the lung and other tissues [20]. Aconitase activity was found to be lower in mitochondria isolated from hyperoxia-exposed AT2 from C57WT, C57MNX and C3HMNX mice (mean [SD], mOD/min/mg protein, 5 [0.04] vs 7 [0.5], 6 [0.02] vs 7 [0.3], and 6 [0.1] vs 6 [0.1] respectively, P < 0.005 for all comparisons) but not in C3HWT mice (7 [0.3] vs 6 [1], P = 0.06) strains. All mice strains had similar AT2 aconitase activity when exposed to normoxia. However oxidative inactivation of aconitase was found to be higher in AT2 from hyperoxia-exposed mice carrying C57 mtDNA (C57WT and C3HMNX) compared to their nuclear counterparts (C57MNX and C3HWT respectively) with C3H mtDNA (**Figure 3B**).

Expression of SIRT3 that modulates mtDNA damage and AT2 apoptosis during oxidative stress was variably modified by hyperoxia exposure in AT2 cells from newborn mice [21]. SIRT3 protein content in AT2 measured by Western blot was reduced by hyperoxia in C57WT and C3HMNX mice (mean [SD], density relative to normoxia-exposed C57WT group, 1.2 [0.01] vs 1 [0.01] and 1 [0.02] vs 4 [0.4] respectively, P < 0.05) but not in C57MNX or C3HWT mice (3 [0.2] vs 4 [0.3] and 3 [0.3] vs 4 [0.4], P = 0.12 and 0.29 respectively). SIRT3 mRNA expression was lower in C57WT, C3HWT and C3HMNX mice (mean [SD], mRNA expression relative to normoxia-exposed C57WT group, 0.7 [0.02] vs 1 [0.1], 1 [0.02] vs 1.7 [0.04], and 0.2 [0.04] vs 2 [0.1] respectively, P < 0.05 for all comparisons) but not in C57MNX mice (1.3 [0.1] vs 1 [0.04], P = 0.1). AT2 SIRT3 protein expression was lower in both normoxia and hyperoxia-exposed C57WT mice compared to C57MNX mice, but only in hyperoxia-exposed C3HMNX mice compared to C3HWT mice (**Figures 3C** and **3E**). However, SIRT3 mRNA expression in AT2 was lower in both normoxia and hyperoxia-exposed mice with C3H mtDNA compared to their nuclear counterparts (C57MNX mice and C3HWT respectively) carrying C57WT (**Figure 3F**). Overall, these findings indicate that mitochondrial oxidative stress in AT2 cells in the lung may vary with differences in the mtDNA carried by these mice.

### 2.4. Hyperoxia-induced oxidative stress in lung homogenates is exacerbated in mice carrying C57 mtDNA

We have previously identified that NMLF from mice carrying C57 mtDNA strains that were exposed to hyperoxia generated more superoxide (O_2_^−^) compared to NMLF from C3H mtDNA strains [16]. To determine whether the higher mtDNA damage and lower aconitase activity that we noted in AT2 from mice with C57 mtDNA compared to mice with C3H mtDNA was associated with globally elevated oxidative stress in their lungs, we measured levels of reactive oxidant species (ROS) such as O_2_^−^ and hydrogen peroxide (H_2_O_2_) as well as markers of lipid and protein oxidation such as malondialdehyde (MDA) and protein carbonyls in mice lung homogenates. Lung O_2_^−^ content was higher in hyperoxia-exposed C57WT and C3HMNX mice (mean [SD], fold change relative to normoxia-exposed C57WT group, 2 [0.3] vs. 1 [0.1] and 4 [0.5] vs. 2 [0.4] respectively, P < 0.01 for both comparisons) but not in C57MNX or C3HWT mice (1.4 [0.1] vs. 1.4 [0.4] and 2.6 [0.5] vs. 2.3 [0.9], P = 0.9 and 0.8 respectively). However, hyperoxia increased lung H_2_O_2_ content in all mice strains (mean [SD], nmol/g of tissue, 56 [9] vs. 25 [4] in C57WT, 31 [5] vs. 21 [4] in C57MNX, 36 [4] vs. 16 [2] in C3HWT and 51 [11] vs. 11 [2] in C3HMNX mice, P < 0.001 for all comparisons). Lung MDA content was higher in hyperoxia-exposed C57WT, C3HWT and C3HMNX mice (mean [SD], nmol/mg of tissue, 3 [0.3] vs. 1.5 [0.3], 2.3 [0.1] vs. 2 [0.2] and 3 [0.1] vs. 2 [0.2] respectively, P < 0.05 for all comparisons) but not in C57MNX mice (2 [0.4] vs. 1.4 [0.4], P = 0.1). However, and similar to lung H_2_O_2_ content, lung protein carbonyl content was noted to be higher in hyperoxia-exposed mice from all strains (mean [SD], nmol/mg of tissue, 8 [1] vs. 2 [0.5] in C57WT, 5 [0.7] vs. 1 [0.4] in C57MNX, 8 [0.8] vs. 4 [0.3] in C3HWT and 11 [2] vs. 5 [1.2] in C3HMNX mice, P < 0.05 for all comparisons).

When exposed to normoxia, lung O_2_^−^ and MDA content was similar in C57WT vs. C57MNX mice and in C3HMNX vs. C3HWT mice whereas lung H_2_O_2_ content was higher in C3HMNX vs. C3HWT mice but similar between C57WT vs. C57MNX mice and lung protein carbonyl content was higher in C57WT vs. C57MNX mice but similar between C3HMNX and C3HWT mice. When exposed to hyperoxia, lung O_2_^−^, H_2_O_2_ and MDA content was higher in C57WT vs. C57MNX mice as well as in C3HMNX vs. C3HWT mice whereas protein carbonyl content was higher only in C57WT compared to C57MNX mice but similar between C3HWT and C3HMNX mice. (**Figures 4A-D**). Taken together, these findings suggest that newborn mice with C57 mtDNA had higher levels of ROS activity in their lungs when exposed to hyperoxia compared to mice with C3H mtDNA.

**Figure 4:**
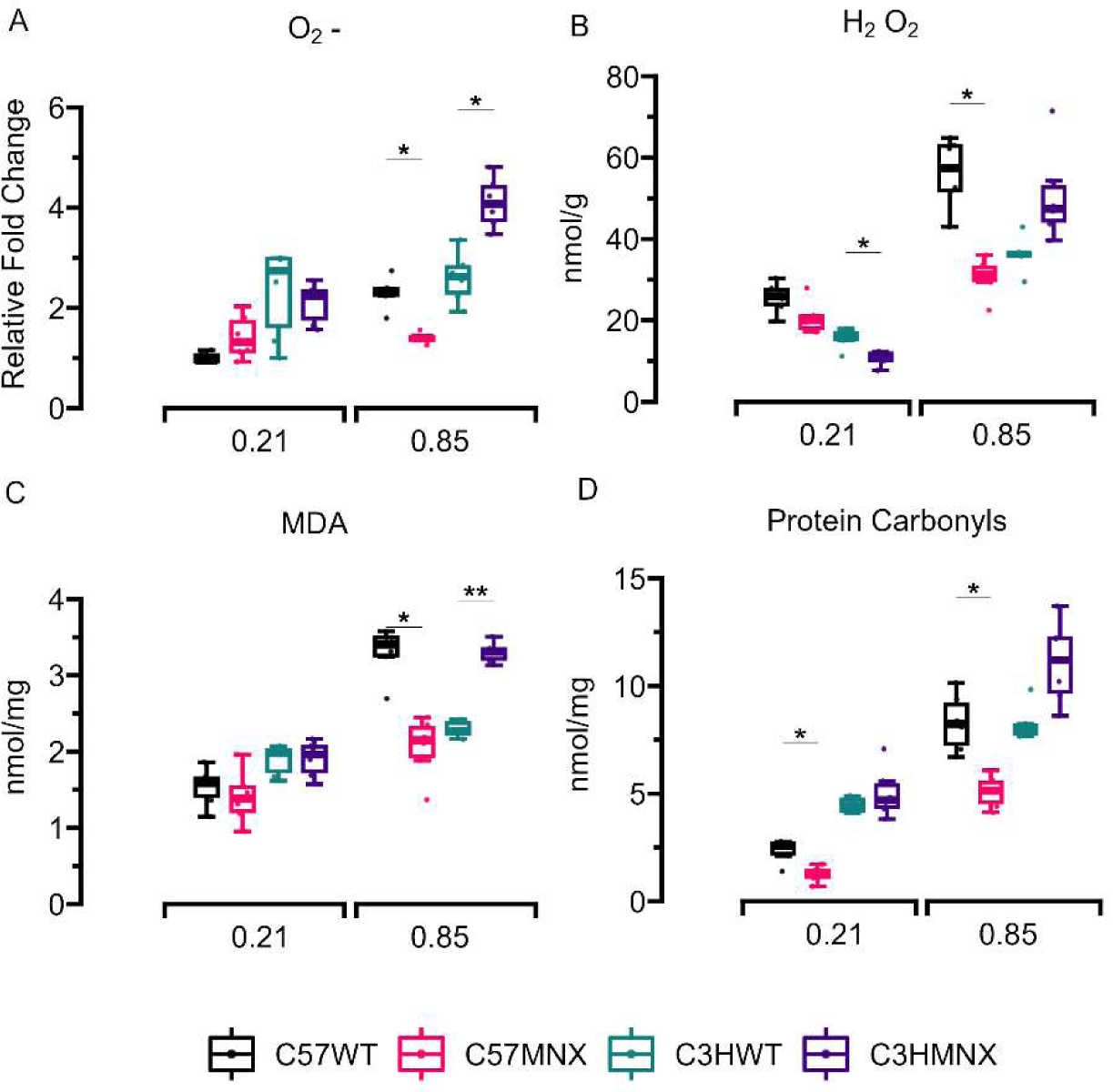
Oxidative stress in whole lung homogenates from mice. **A**: O_2_^−^. **B**: H_2_O_2_. **C**: MDA. **D**: Protein carbonyls. N = minimum of 6 mice/group. All data were analyzed by 2-way ANOVA or Kruskal-Wallis tests, followed by post hoc analyses. Box represents median/interquartile range, whiskers represent maximum and minimum values. * and ** represent p-values < 0.05 and < 0.005 respectively.

### 2.5. Lung transcriptome may be differentially modified by hyperoxia in mice with different mtDNA

Mito-nuclear interactions modify gene expression and the MNX model we used in our previous and current studies have been used to evaluate the independent effects of mtDNA variations on gene expression in several tissues and disease models [22,23]. Having previously found that mtDNA variations modify hyperoxic lung injury severity in newborn mice we investigated gene expression in the lungs of both WT and MNX mice exposed to normoxia or hyperoxia in this study [16]. Principal component analysis (PCA) showed that samples from mice with similar nuclear genetic backgrounds and exposures (normoxia vs. hyperoxia) clustered together. However, we also noted a higher degree of separation between samples from mice with similar nuclear backgrounds but different mitochondrial backgrounds during hyperoxia as compared to normoxia (**Figure 5A**). Interestingly, more genes were differentially regulated by hyperoxia (genes with LFC in expression > 2 and FDR p-value < 0.05 in hyperoxia-exposed mice compared to normoxia-exposed mice) in mice carrying C57 mtDNA (490 up/634 down regulated genes in C57WT mice and 474 up/621 down regulated genes in C3HMNX mice) compared to mice carrying C3H mtDNA (386 up/493 down regulated genes in C57MNX mice and 349 up/439 down regulated genes in C3HWT mice). We also noted that 50 genes were commonly differentially regulated (33 upregulated/17 downregulated) in mice strains carrying C57 mtDNA (C57WT and C3HMNX) whereas 19 genes were commonly differentially regulated by hyperoxia (10 upregulated/9 downregulated) in mice strains with C3H mtDNA (C57 mtDNA and C3H mtDNA) as shown in **Figures 5B-C**.

**Figure 5:**
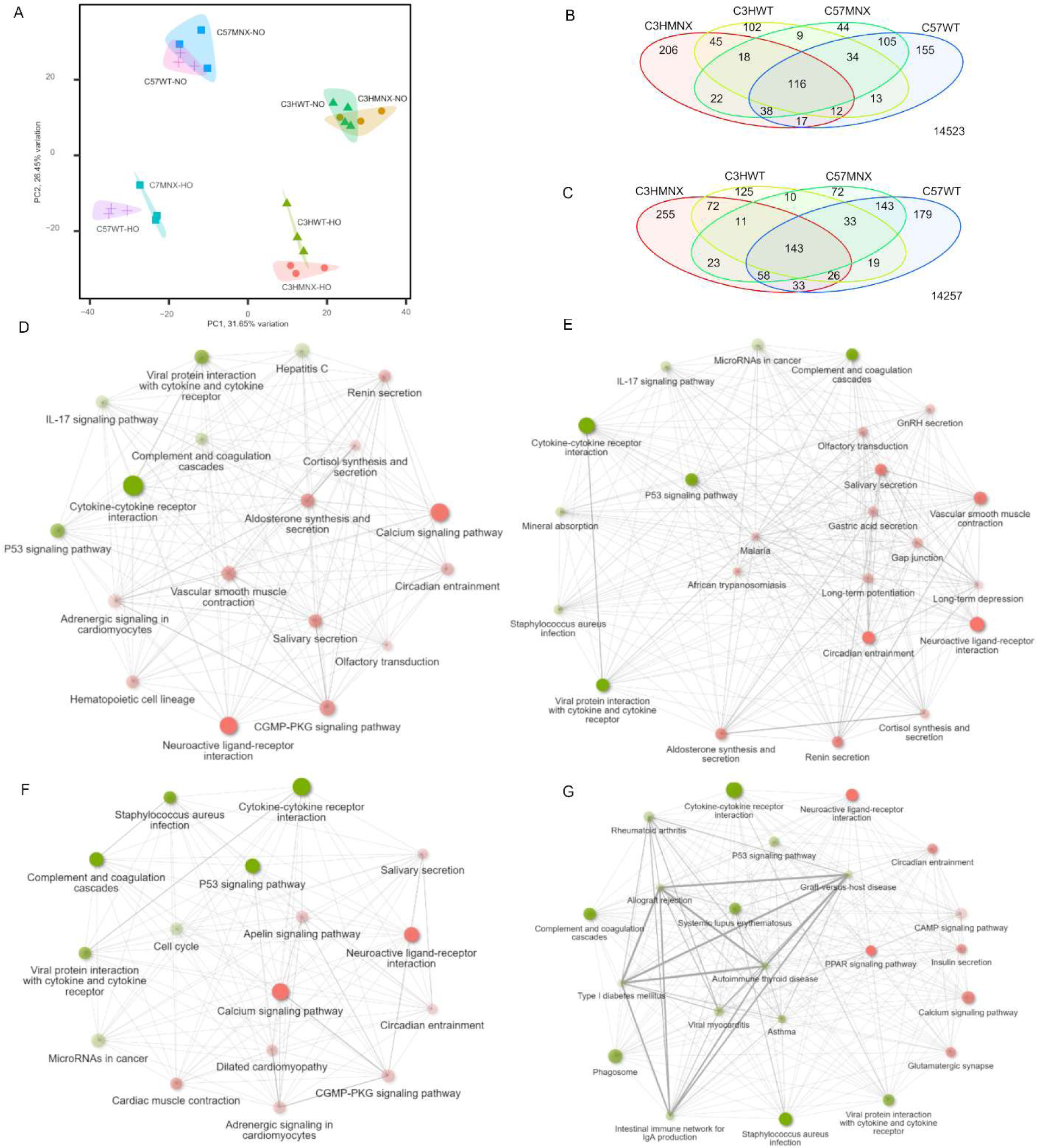
Hyperoxia-induced differences in lung transcriptomes of WT and MNX mice. **A**: Principal component analysis of RNA sequencing data indicating that clustering of lung transcriptomes based on nuclear backgrounds as well as a clearer separation between groups exposed to hyperoxia compared to normoxia. **B**: Venn Diagrams of differentially Downregulated and **C**: upregulated genes showing overlaps between groups. Enrichment maps of ORA for KEGG pathways are shown for (**D**) C57WT (**E**) C57MNX (**F**) C3HWT and (**G**) C3HMNX mice.

Over-representation analysis (ORA) of gene sets differentially regulated by hyperoxia exposure (LFC > 1 and FDR p-value < 0.05) from each mice strain for enriched pathways (FDR p-value < 0.05) from the Kyoto Encyclopedia of Genes and Genomes (KEGG) compendium identified that pro-inflammatory pathways such as *Viral protein interaction with cytokine and cytokine receptor*, *Cytokine-cytokine receptor interaction* and *Complement and coagulation cascade*, as well as *p53 signaling* which is known to protect against inflammatory and oxidant injury were enriched by hyperoxia in the lungs of all mice strains (**Figures 5D-G, Figure 6** and **Tables 1** to **4**) ORA also identified that hyperoxia upregulated the *Hepatitis C* pathway which is associated with cell cycle arrest in C57 WT mice and the *Mineral absorption* pathway which plays roles in surfactant and redox metabolism in C57MNX mice [24,25]. Furthermore, pathways related to glutamatergic signaling (*long-term potentiation* and *glutamatergic synapse*) which is known to have pro-fibrotic effects in the developing lung were inhibited during hyperoxia exposure in the lungs of C57MNX mice [26]. Another interesting observation was that pathways related to the cell cycle were upregulated whereas Apelin signaling, which inhibits branching morphogenesis was downregulated in C3HWT mice exposed to hyperoxia [27]. We also noted more pronounced upregulation of other pro-inflammatory pathways (*Asthma, Tuberculosis, Systemic lupus erythematosus, Rheumatoid arthritis,* and *Viral myocarditis)* as well as downregulation of pathways associated with anti-inflammatory and anti-fibrotic effects in the developing lung such as the PPAR signaling pathway in hyperoxia-exposed C3HMNX mice but not in the lungs of C3HWT mice [28].

**Figure 6:**
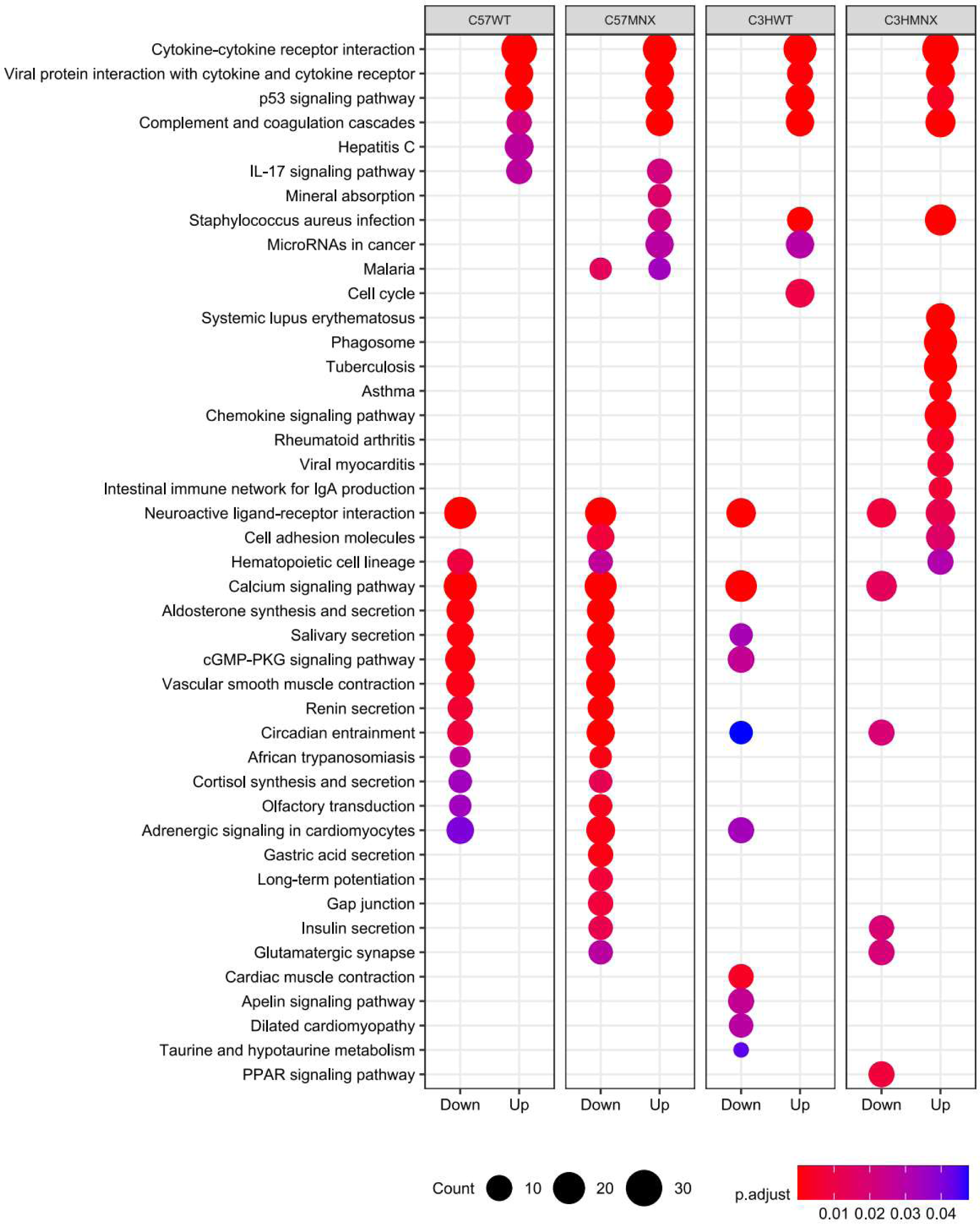
Dotplots of ORA for KEGG pathways in the lung transcriptomes of mice. The top 15 pathways were identified using gene lists filtered from differential expression results (LFC > 1 and FDR p-value < 0.05) that were analyzed using ClusterProfiler and R. Count represents number of genes that were enriched in each pathway.

**Table 1:** GSEA analysis of hyperoxia regulated genes from C57WT mice lungs

| Pathway | t-statistic | FDR P-value | Number of Genes |
| --- | --- | --- | --- |
| Cytokine-cytokine receptor interaction | 5.05 | 7.3e-05 | 164 |
| Neuroactive ligand-receptor interaction | 5.04 | 7.3e-05 | 130 |
| Viral protein interaction with cytokine and cytokine receptor | 4.24 | 2.9e-03 | 58 |
| Spliceosome | -5.3 | 6.9e-05 | 123 |
| Ribosome | -5.1 | 6.9e-05 | 123 |
| Ubiquitin mediated proteolysis | -4.3 | 1.6e-03 | 134 |
| Protein processing in endoplasmic reticulum | -3.84 | 5.1e-03 | 155 |
| MRNA surveillance pathway | -3.77 | 7.2e-03 | 84 |
| **Mitophagy** | -3.29 | 2.8e-02 | 65 |
| Autophagy | -3.28 | 2.7e-02 | 131 |
| Amyotrophic lateral sclerosis | -3.09 | 3.7e-02 | 328 |

**Table 2:**
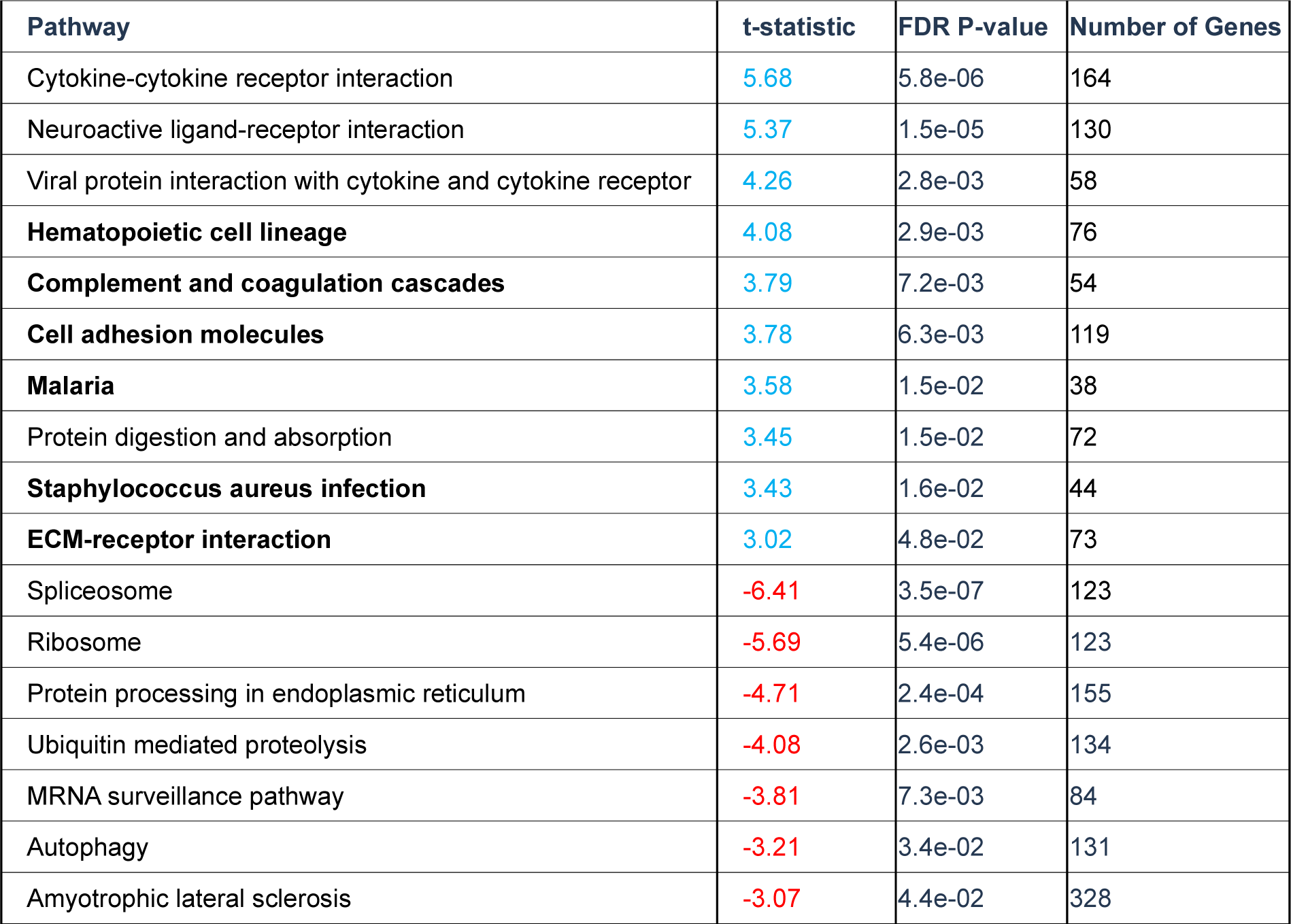
GSEA analysis of hyperoxia regulated genes from C57MNX mice lungs

| Pathway | t-statistic | FDR P-value | Number of Genes |
| --- | --- | --- | --- |
| Cytokine-cytokine receptor interaction | 5.68 | 5.8e-06 | 164 |
| Neuroactive ligand-receptor interaction | 5.37 | 1.5e-05 | 130 |
| Viral protein interaction with cytokine and cytokine receptor | 4.26 | 2.8e-03 | 58 |
| **Hematopoietic cell lineage** | 4.08 | 2.9e-03 | 76 |
| **Complement and coagulation cascades** | 3.79 | 7.2e-03 | 54 |
| **Cell adhesion molecules** | 3.78 | 6.3e-03 | 119 |
| **Malaria** | 3.58 | 1.5e-02 | 38 |
| Protein digestion and absorption | 3.45 | 1.5e-02 | 72 |
| **Staphylococcus aureus infection** | 3.43 | 1.6e-02 | 44 |
| **ECM-receptor interaction** | 3.02 | 4.8e-02 | 73 |
| Spliceosome | -6.41 | 3.5e-07 | 123 |
| Ribosome | -5.69 | 5.4e-06 | 123 |
| Protein processing in endoplasmic reticulum | -4.71 | 2.4e-04 | 155 |
| Ubiquitin mediated proteolysis | -4.08 | 2.6e-03 | 134 |
| MRNA surveillance pathway | -3.81 | 7.3e-03 | 84 |
| Autophagy | -3.21 | 3.4e-02 | 131 |
| Amyotrophic lateral sclerosis | -3.07 | 4.4e-02 | 328 |

**Table 3:** GSEA analysis of hyperoxia regulated genes from C3HWT mice lungs

| Pathway | LFC | FDR P-value | Number of Genes |
| --- | --- | --- | --- |
| Cytokine-cytokine receptor interaction | 5.27 | 4.6e-05 | 164 |
| Viral protein interaction with cytokine and cytokine receptor | 4.31 | 3.4e-03 | 58 |
| Complement and coagulation cascades | 4.18 | 3.4e-03 | 54 |
| **P53 signaling pathway** | 3.56 | 2.2e-02 | 67 |
| Neuroactive ligand-receptor interaction | 3.41 | 2.5e-02 | 130 |
| Ribosome | -4.55 | 8.5e-04 | 123 |
| Amyotrophic lateral sclerosis | -4.47 | 8.5e-04 | 328 |
| Spliceosome | -4.37 | 8.5e-04 | 123 |
| **Huntington disease** | -4.29 | 8.5e-04 | 274 |
| MRNA surveillance pathway | -3.77 | 8.3e-03 | 84 |
| Autophagy | -3.45 | 1.8e-02 | 131 |
| Protein processing in endoplasmic reticulum | -3.27 | 2.8e-02 | 155 |
| Ubiquitin mediated proteolysis | -3.19 | 3.2e-02 | 134 |

**Table 4:**
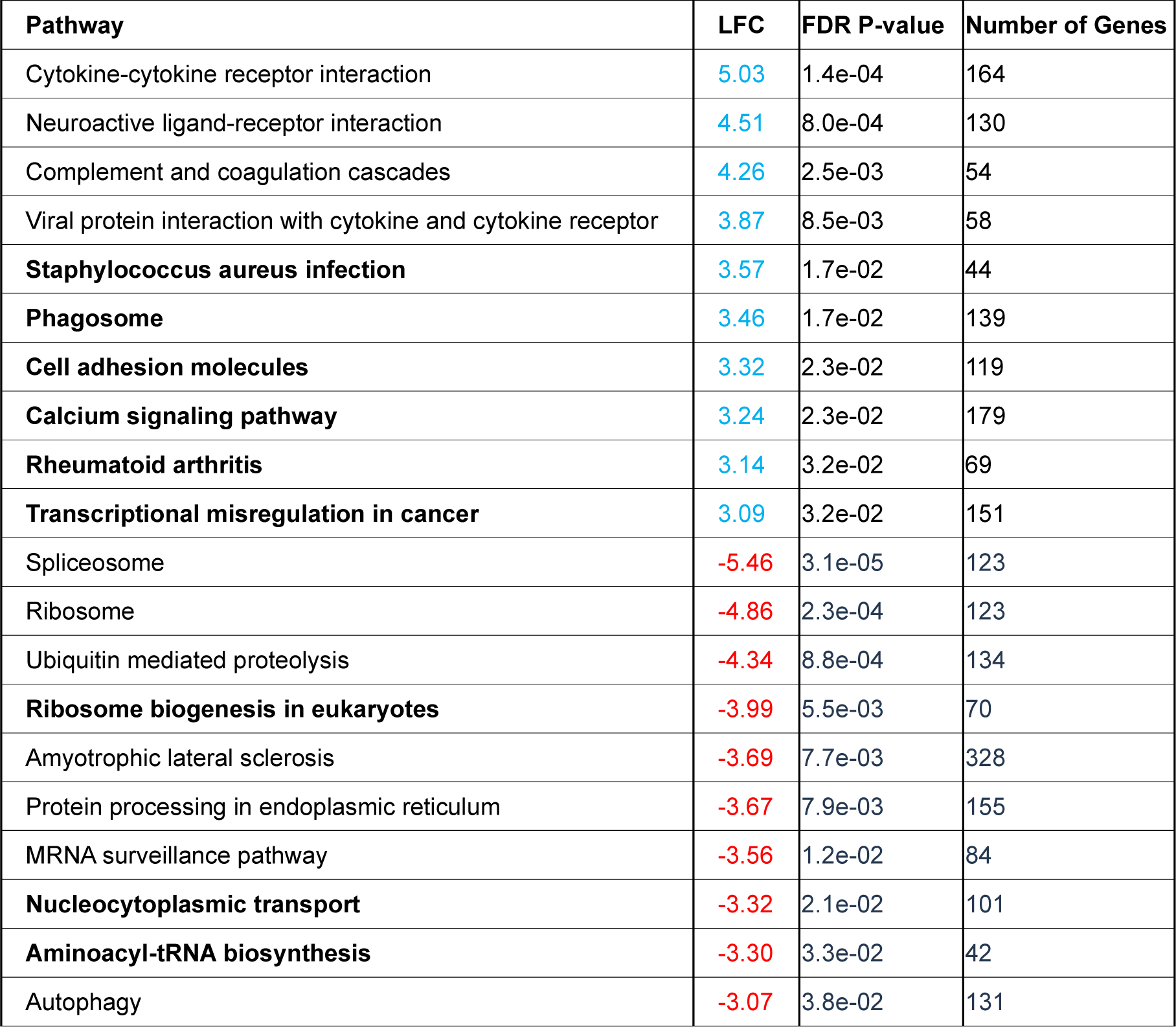
GSEA analysis of hyperoxia regulated genes from C3HMNX mice lungs

We also performed gene set enrichment analysis (GSEA) of ranked lists of all genes expressed in mice lungs using the threshold-free Generally Applicable Gene-set Enrichment (GAGE) method to detect additional KEGG pathways with small but coordinated changes that were significantly (LFC > 1 and FDR < 0.05) differentially modified by hyperoxia [29,30]. While hyperoxia-induced changes in lung transcriptome largely produced similar effects on KEGG pathways in C57WT vs. C57MNX and in C3HMNX vs. C3HWT mice (such as downregulation of the Autophagy pathway), we also noted differences such as inhibition of the *Mitophagy* pathway in C57WT mice but not in C57MNX mice (primarily related to *PINK1* inhibition), as well as of inhibition of pathways related to pulmonary vascular remodeling (*Nucleocytoplasmic transport*) and protein synthesis (*Aminoacyl-tRNA synthesis*) in C3HMNX but not C3HWT mice lungs. These observations indicate that hyperoxia-induced changes in the lung transcriptome such as upregulation of inflammatory pathways and downregulation of protective/repair mechanisms such as autophagy may vary with differences in mtDNA in newborn mice (**Tables 1-4, Figure 7**) [31].

**Figure 7:**
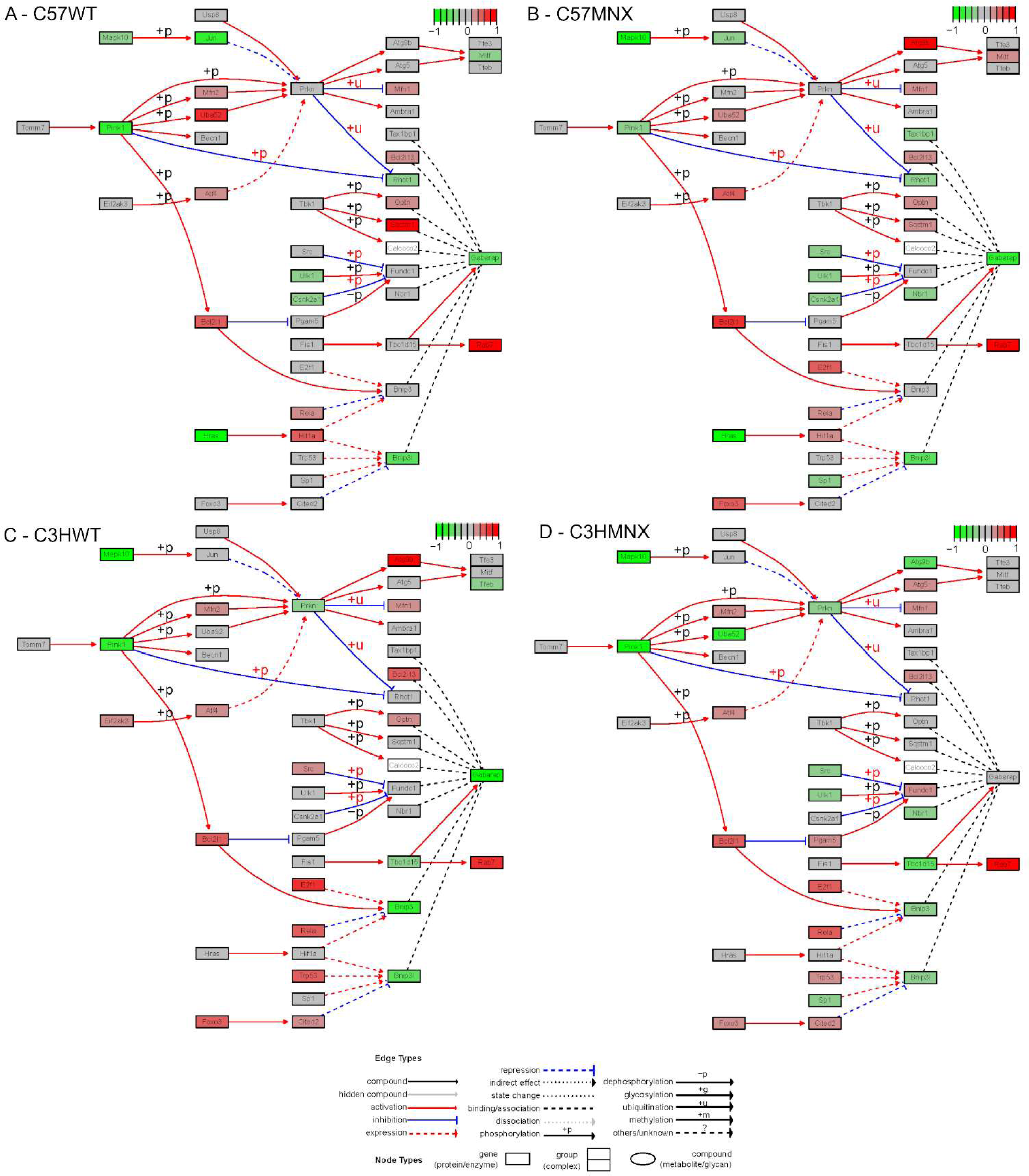
KEGG map for the Mitophagy in Animals pathway in the lung transcriptomes of WT and MNX mice. Map was created using the Graphviz package and R and shows interrelationships between constituent genes with differentially regulated genes shaded as green (down) or red (up). **A**: C57WT, **B**: C57MNX, **C**: C3HWT and **D**: C3HMNX mice. PINK1 was downregulated to a greater degree by hyperoxia in C57MWT mice compared to C57MNX mice.

### 2.6. Inflammatory cytokines are increased in the lungs of hyperoxia-exposed mice with C57 mtDNA

In addition to pulmonary bioenergetics and oxidant stress in the lungs of hyperoxia-exposed newborn mice with C57 mtDNA and mice with C3H mtDNA we also analyzed cytokines in the lungs of these mice by ELISA to further explore the findings that hyperoxia exposure upregulated several pathways related to cytokine and chemokine activity in the lung transcriptomes of newborn mice. As shown in **Figure 8**, hyperoxia increased the pro-inflammatory cytokines IL1-β and IL-6 in all mice strains (mean [SD], log_10_ pg/mL, IL1-β: 3 [0.1] vs. 2.7 [0.02], 3 [0.02] vs. 2.4 [0.1], 3 [0.04] vs. 2.42 [0.04] and 3 [0.08] vs. 2.49 [0.03] mice, IL-6: 4 [0.1] vs. 3 [0.02], 4 [0.1] vs. 3 [0.02] in, 3 [0.3] vs. 3 [0.01] in and4 [0.3] vs. 3 [0.01] in C57WT, C57MNX, C3HWT and C3HMNX mice respectively, P < 0.005 for all comparisons). TNF-α and IFN-γ were increased by hyperoxia only in C57WT and C3HMNX mice (TNF-α: 4 [0.2] vs. 3 [0.01] and 4 [0.06] vs. 3 [0.02], IFN-γ: 2.9 [0.02] vs. 2.6 [0.03] and 2.5 [0.02] vs. 2 [0.03] respectively, P < 0.005 for all comparisons) but not in C57MNX or C3HWT mice (TNF-α: 3 [0.1] vs. 3 [0.02] and 3 [0.1] vs. 3 [0.04], P = 0.8 and 0.2; IFN-γ: 3 [0.03] vs. 2.5 [0.08] and 2.4 [0.05] vs. 2 [0.06]), P = 0.5 and 0.6 respectively). Notably, levels of the neutrophil chemoattractant CXCL1 were higher in the lungs of hyperoxia-exposed C57WT, C57MNX and C3HWT mice (3.4 [0.04] vs. 3 [0.01], 3 [0.04] vs. 2 [0.02], and 2.94[0.01] vs. 2.91[0.02] respectively, P < 0.05 for all comparisons) but lower in hyperoxia-exposed C3HMNX mice (2.8 [0.1] vs. 3 [0.01], P < 0.001).

**Figure 8:**
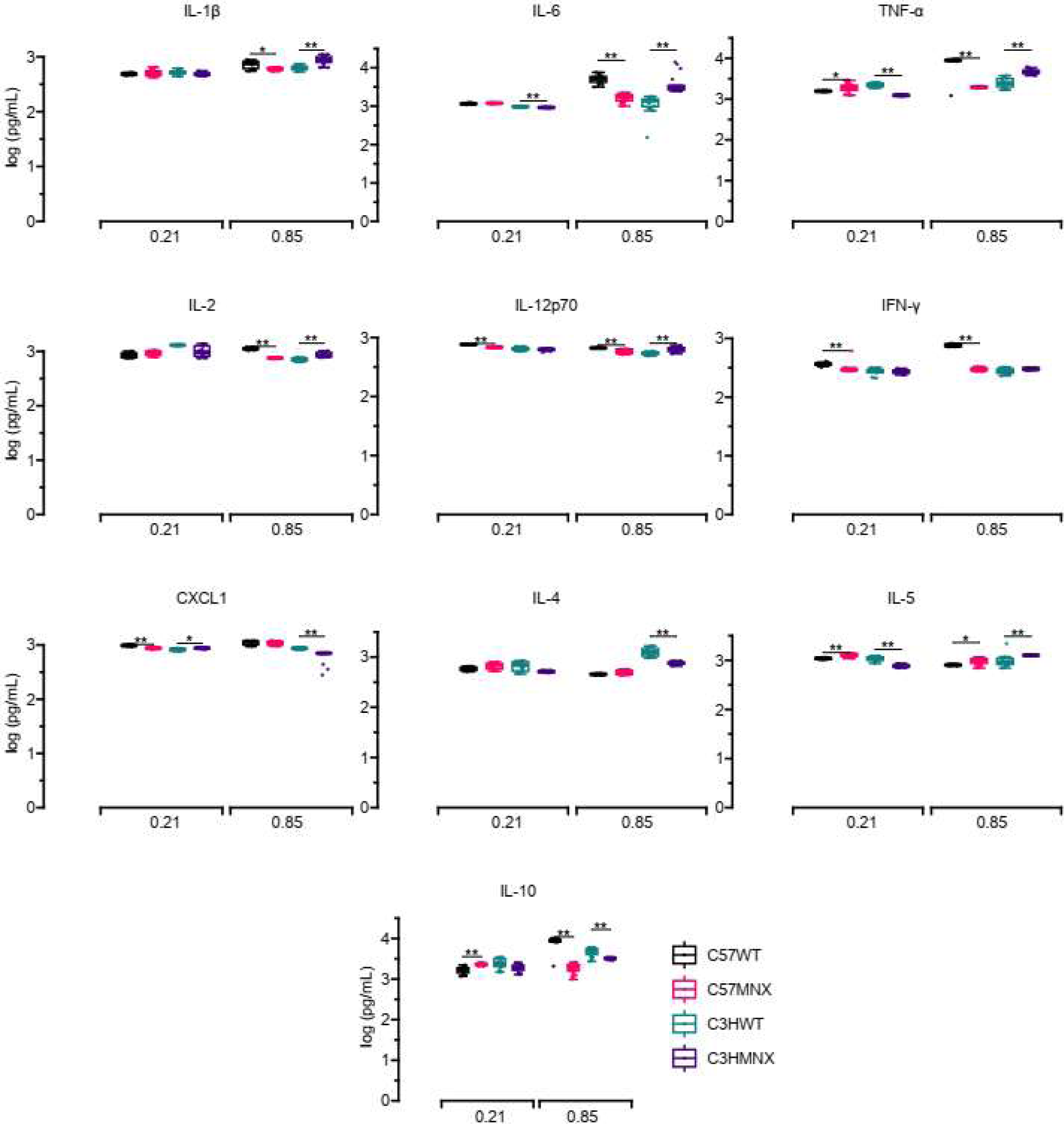
Inflammatory cytokine content in the lungs of MNX and WT mice. Plots depict levels of cytokines expressed as log-transformed picograms/mL of lung tissue lysate for IL-1β, IL-6, TNF-α, IL-2, IL-12p70, IFN-γ, KC/GRO (CXCL1), IL-4, IL-5, and IL-10. N = minimum of 12 mice/group. All data were analyzed by 2-way ANOVA or Kruskal-Wallis tests, followed by post hoc analyses. Box represents median/interquartile range, whiskers represent maximum and minimum values. * and ** represent p-values < 0.05 and < 0.005 respectively.

When exposed to hyperoxia, lungs from mice with C57 mtDNA (C57WT and C3HMNX) had higher levels of BPD-associated cytokines such as IL1-β, IL-6, and TNF-α (as well as IL-2, which has been proposed to have a beneficial effect in fetal lung injury models and Il-12 whose role in BPD remains unclear) compared to mice with C3H mtDNA (C57MNX and C3HWT) respectively [32,33]. Levels of IFN-γ were higher in lungs from hyperoxia-exposed C57WT mice compared to C57MNX mice, but similar in C3HMNX vs. C3HWT mice, whereas CXCL1 was similar between hyperoxia-exposed C57WT vs. C57MNX mice, but lower in C3HMNX mice compared to C3HWT mice. Additionally, IL-4, which is known to be anti-inflammatory but may not have a pathogenic role in BPD, was similar between C57WT vs. C57MNX mice but lower in C3HMNX mice compared to C3HWT mice, whereas levels of the eosinophil regulatory cytokines IL-5 and IL-10 were found to be lower in both MNX strains compared to their WT counterparts [34]. These results indicate that hyperoxia-induced cytokine profiles in mice lungs vary with mtDNA differences, and lungs of mice carrying C57 mtDNA have a higher level of pro-inflammatory cytokine activity compared to mice carrying C3H mtDNA (**Figure 8**).

### 2.7. Hyperoxia-induced decrease in autophagy and mitophagy is higher in AT2 and NMLF from mice with C57 mtDNA

Since GSEA of the lung transcriptome in mice found a global downregulation of autophagy and differential modulation of mitophagy in C57WT vs. C57MNX mice, we investigated these pathways in mouse lungs. Transmission electron microscopy (TEM) of lung sections from normoxia and hyperoxia-exposed mice showed that lung sections from hyperoxia-exposed C57WT, C3HWT and C3HMNX mice had markedly fewer autophagic vacuoles per cell compared to their controls (mean [SD], vacuoles/cell relative to normoxic C57WT mice, 0.3 [0.2] vs. 1 [0.2], 1 [0.1] vs. 1.3 [0.1] and, 0.4 [0.1] vs. 0.9 [0.1] respectively, P < 0.05 for all comparisons) but not in C57MNX mice (0.9 [0.2] vs. 1.4 [0.4], P = 0.2). Unexpectedly, there were no differences in relative mitochondrial area per cell between hyperoxia and normoxia-exposed mice from each strain, nor between strains during either normoxia or hyperoxia, which suggests that mitophagy may not be differentially regulated by hyperoxia or by mtDNA variations.(**Figures 9 A-C**).

**Figure 9:**
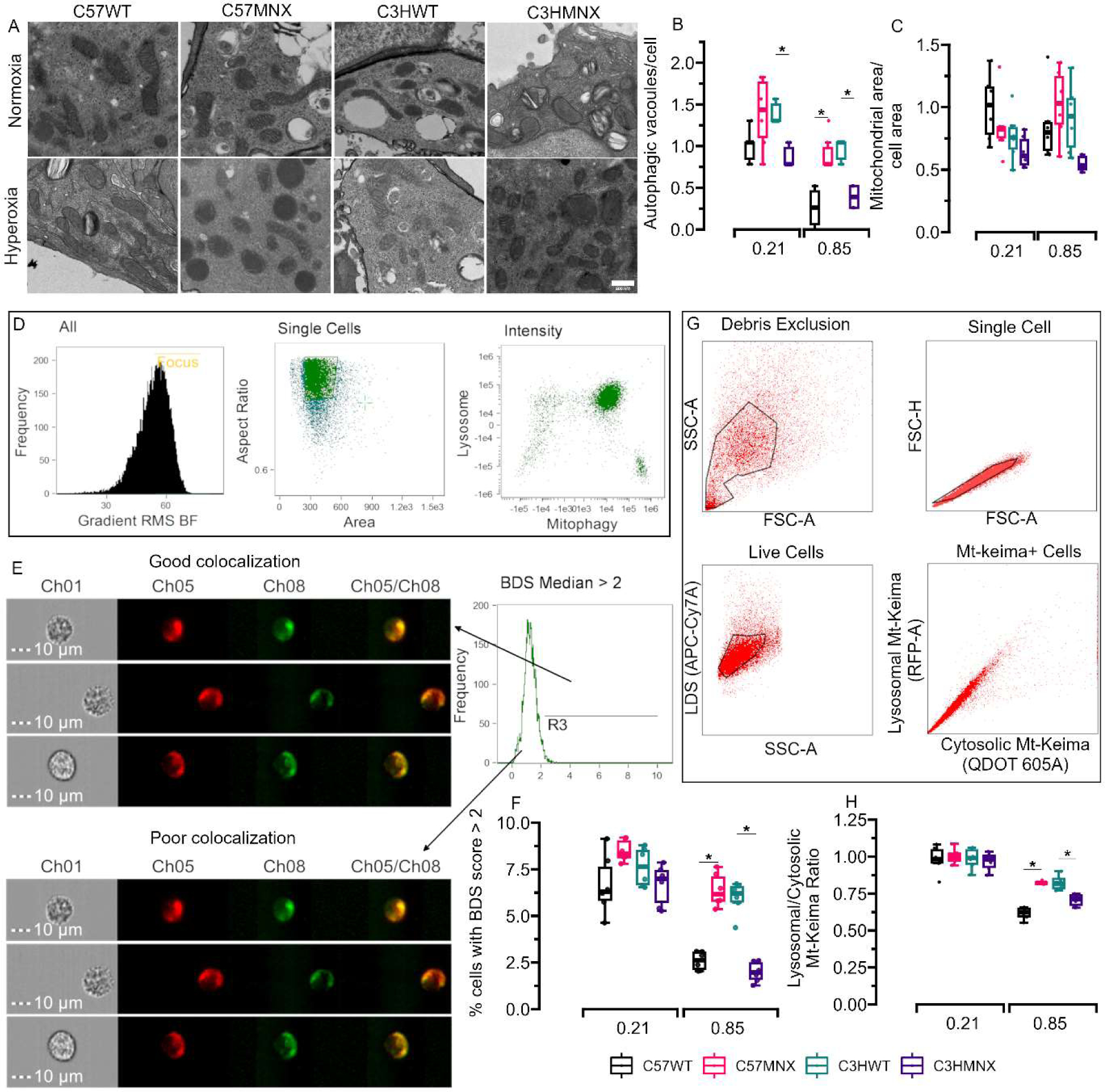
Autophagy and mitophagy in the lungs of MNX and WT mice. (**A**) Representative TEM images of lung sections showing more autophagy in normoxic mice compared to hyperoxic mice in all groups. (**B**) Autophagic vacuoles per cell in TEM images. (**C**) Mitochondrial area/per cell. (**D-E**) Representative imaging flow cytometry workflow for gating and measuring BDS scores using co-localization of mitochondrial dye and lysosomal dye in AT2 cells. (**F**) BDS scores in AT2 cells. (**G**) Representative flow cytometry workflow for gating NMLF to measure mitophagy using mt-Keima fluorescence measurements in mitochondrial and lysosomal compartments. (**H**) Lysosomal/mitochondrial mt-Keima fluorescence ratio in NMLF. N = minimum of 3 mice/group. All data were analyzed by 2-way ANOVA or Kruskal-Wallis tests, followed by post hoc analyses. Box represents median/interquartile range, whiskers represent maximum and minimum values. * and ** represent p-values < 0.05 and < 0.005 respectively.

However, mitochondrial quality control through mitophagy has been found to play a prominent role in various lung pathologies and we have previously shown that MSC from infants with BPD had lower mitophagy compared to MSC from infants without BPD. Therefore, we also investigated whether hyperoxia induced differential regulation of mitophagy is modulated by mtDNA variations through mitochondria-lysosomal dye colocalization to calculate bright-detail similarity (BDS) scores using imaging flow cytometry in AT2 cells and a transfection-based approach to measure lysosomal/cytosolic (mitochondrial) fluorescence of mt-Keima in NMLF obtained from the lungs of these mice (**Figures 9D-E, G**). BDS scores of AT2 C57WT, C57MNX and C3HMNX mice exposed to hyperoxia were lower (indicative of decreased mitophagy) compared to controls (mean [SD], fraction of cells with BDS scores > 2, 3 [0.5] vs. 7 [1.6], 6 [0.9] vs. 9 [0.6] and 2 [0.5] vs. 7 [1.1] respectively, P < 0.05 for all comparisons) but similar in C3HWT mice (6 [0.9] vs. 8 [1.03], P =0.06). Mice with C57 mtDNA (C57WT and C3HMNX) had lower AT2 BDS scores compared to mice with C3H mtDNA (C57MNX and C3HWT) mice respectively (**Figure 9F**). Similarly, NMLF lysosomal/cytosolic mt-Keima fluorescence was lower (indicative of decreased mitophagy) in hyperoxia vs. normoxia-exposed mice from all mice strains (mean [SD], fluorescence ratio relative to normoxia-exposed C57WT mice, 0.8 [0.01] vs. 1 [0.05] in C57WT, 0.6 [0.04] vs. 1 [0.09] in C57MNX, 0.8 [0.05] vs. 11[0.07] in C57WT and 0.7 [0.04] vs. 1 [0.06] in C57 MNX strains, P < 0.05 for all comparisons. NMLF from hyperoxia-exposed mice with C57 mtDNA (C57WT and C3HMNX) mice were also noted to have lower lysosomal mt-Keima fluorescence compared to mice with C3H mtDNA (C57MNX and C3HWT respectively), indicating that mitophagy in lung cells may be modulated by mtDNA variations in hyperoxic neonatal lung injury (**Figure 9H**).

### 2.8. MtDNA modulates hyperoxia-induced decrease in AT2 Pink1, Parkin and Pgc1-α expression

Western blot analysis of mitophagy-related proteins PINK1 and Parkin showed a hyperoxia-induced reduction in PINK1 protein content in AT2 cells from C57WT, C3HWT and C3HMNX strains but reduced parkin protein content in AT2 cells from C3HWT and C3HMNX strains only. Additionally, qPCR analysis in AT2 cell lysates found hyperoxia-induced reduction in Pink1 (supporting the changes noted in the KEGG mitophagy pathway analysis in **Figure 7**) and Parkin mRNA in AT2 cells from all 4 mice strains. PINK1 and Parkin protein and mRNA expression in AT2 were lower in both C57 mtDNA-carrying strains (C57WT and C3HMNX) compared to their C3H mtDNA carrying nuclear counterparts (C57MNX and C3HWT respectively), providing further evidence for our hypothesis that hyperoxia-mediated decrease in lung mitophagy may be modulated by may be modulated by mtDNA variations and differences in mitochondrial function (**Figures 10A-B**).

**Figure 10:**
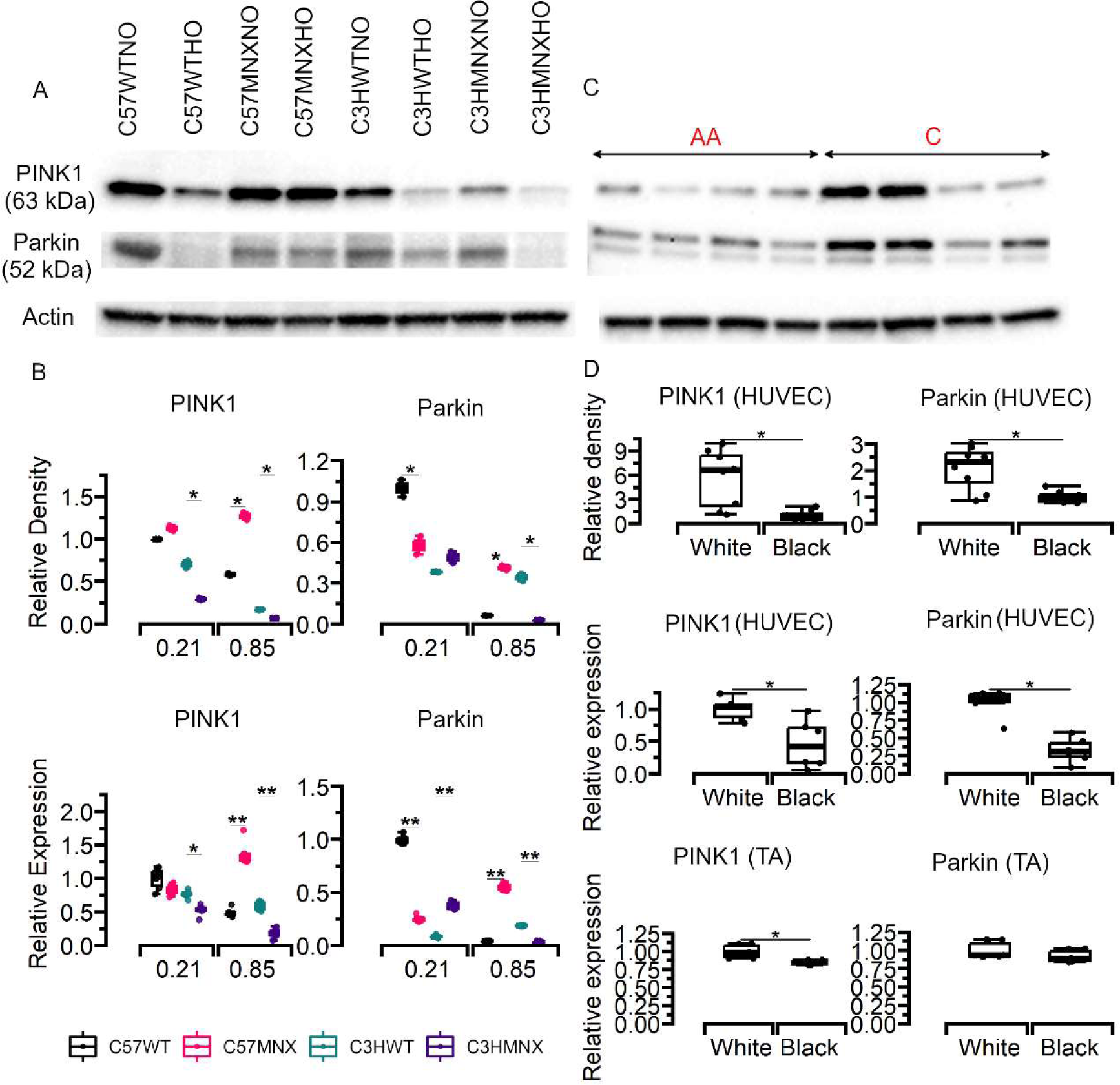
PINK1 and Parkin expression in AT2 from mice and HUVEC and tracheal aspirates from infants. (**A**) Representative Western blot for PINK1 and Parkin protein expression in AT2 cells. qPCR. **(B)** Densitometry and relative expression of Pink1 and Parkin from AT2 cells. **(C)** Representative Western blot for PINK1 and Parkin protein expression in HUVEC from Black and White infants who later developed BPD. (**D**) Densitometry and relative mRNA expression of PINK1 and Parkin in HUVEC and tracheal aspirates obtained around 28 days of life from infants who developed BPD. N = 6 mice/group and 8 infants per group. All data were analyzed by 2-way ANOVA or Kruskal-Wallis tests, followed by post hoc analyses. Box represents median/interquartile range, whiskers represent maximum and minimum values. * and ** represent p-values < 0.05 and < 0.005 respectively.

In addition to measuring mitochondrial surface area in TEM lung sections, we also measured expression of ETC complex protein subunits to identify if changes in mitophagy were also associated with significant changes in mitochondrial content in AT2 cells. Similar to the findings noted in the TEM sections, no significant differences in ETC subunit protein content were noted between hyperoxia and normoxia-exposed mice from each strain, or between strains during either normoxia or hyperoxia (**Figures 11A-B**). However, since hyperoxia reduced PPAR signaling (which is a master regulator of mitochondrial biogenesis) in the lung transcriptome of C3HMNX mice, we measured *Pgc1-α* mRNA in AT2 cells through qPCR and found that while hyperoxia decreased *Pgc1-α* expression in AT2 from all strains, mice with C57 mtDNA (C57WT mice and C3HMNX) had lower AT2 *Pgc1-α* expression with hyperoxia compared to mice with C3H mtDNA (C57MNX and C3HWT respectively) [35]. These results indicate that relatively higher levels of mitochondrial biogenesis may contribute to the similarity in mitochondrial content between the various mice strains despite the differences in mitophagy that were noted in the lungs of these mice (**Figure 11C**).

**Figure 11:**
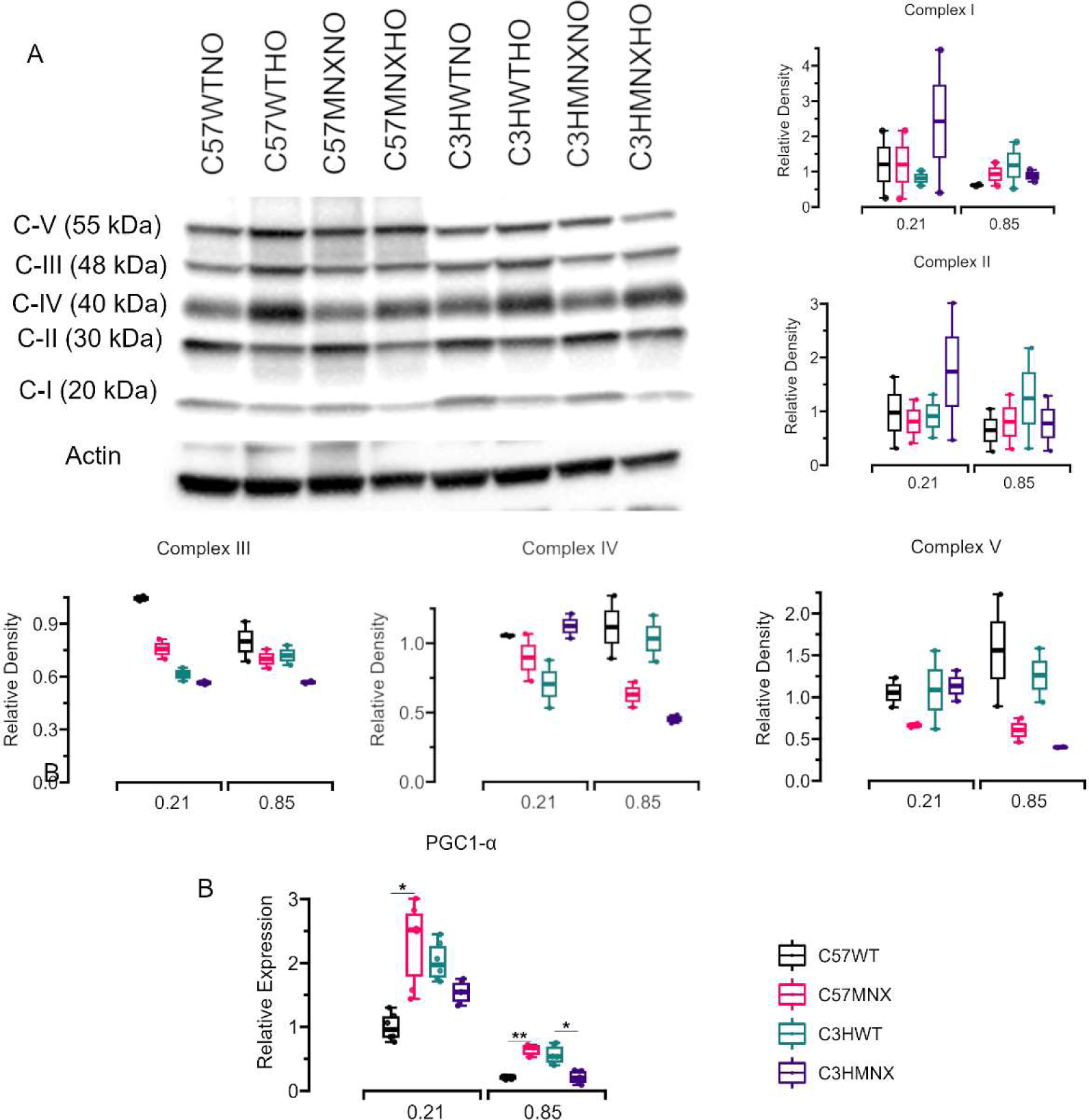
ETC subunit and PGC1-α expression in AT2 cells from mice. (**A**) Representative Western blots for NDUFB8 (Complex I), SDHB (Complex II), UQCR2 (Complex III), MTCO1 (Complex IV) and ATP5A (Complex V) protein expression in AT2 cells. (**B**) PGC1-α mRNA expression in AT2 cells from mice. N = Minimum of 3 mice/group. All data were analyzed by 2-way ANOVA or Kruskal-Wallis tests, followed by post hoc analyses. Box represents median/interquartile range, whiskers represent maximum and minimum values. * and ** represent p-values < 0.05 and < 0.005 respectively.

### 2.9. PINK1 and SIRT3 expression is lower in Black ELBW infants with BPD compared to White infants with BPD

In addition to investigating possible mechanisms associated with mitochondrial function that may have a role in mediating differences in lung injury severity in mice with different mtDNA haplogroups, we also used HUVEC and tracheal aspirates (TA) obtained from a cohort of Black and White infants (self-reported race) who developed BPD (n = 8 per group) to measure SIRT3 as a marker for mitochondrial oxidant stress and PINK1 and Parkin expression as markers for mitophagy. Western blot analysis found that Black infants with BPD had decreased HUVEC SIRT3, PINK1 and Parkin protein and mRNA expression at birth as well as decreased SIRT and PINK1 mRNA but similar Parkin mRNA expression in their tracheal aspirates at 28 days of life (**Figures 3C-D, G-I and 10C-D**). These results suggest that mitophagy and mitochondrial oxidant stress may vary between infants with different ethnicities with similar risk for BPD.

## 3. Discussion

Alveolar formation has been found to require differential activity of mitochondria in various progenitor lung cells, and mitochondrial function is being increasingly recognized as a key contributor to lung diseases such as BPD in which oxidative stress plays an important role [36,37]. Our previous work identified that mice with C57 mtDNA had more severe lung injury, decreased alveolarization and fibroblast bioenergetic dysfunction when exposed to hyperoxia compared to mice carrying C3H mtDNA [16]. In this study we identified increased hyperoxia-induced bioenergetic dysfunction in alveolar epithelial cells from mice with C57 mtDNA when compared to mice with C3H mtDNA. Cellular bioenergetics and ATP availability influence cell function and survival, and differences in bioenergetic function have been associated with biologic variability in maintenance of health and predisposition to diseases [38,39]. Lung development requires substantial energy and alveolar epithelial cells rely on mitochondria-derived ATP to continuously produce and release surfactant, and bioenergetic function may have a profound impact on alveolar epithelial cell survival as well as on associated lung pathologies such as COPD, PH and IPF [37,40–42]. MtDNA-encoded proteins are essential for assembly of competent respiratory chains, and previous studies have found that oxidative mtDNA mutations as well as inherited mtDNA variations lead to differences in mitochondrial bioenergetic function [43,44]. Therefore, our findings that mtDNA variation is associated with differences in the severity of neonatal hyperoxic lung injury as well as alveolar epithelial cell mitochondrial function suggest that predisposition to BPD may be modified by inherited (as well as acquired) mtDNA variants in infants.

Reduced ΔΨm is a hallmark of mitochondrial dysfunction as ETC bioenergetic and redox function are crucial for the maintenance of inner mitochondrial membrane potential and ΔΨm is a major driving force for mitochondrial ATP synthesis [45]. Opening of the MPTP is considered to be a critical factor that leads to the collapse of the ΔΨm, release of cytochrome C into the cytosol and subsequent initiation of apoptotic mechanisms [46]. Heterogeneity of ΔΨm has been observed in association with differences in OXPHOS capacity in mitochondrial cybrids carrying different mtDNA haplogroups [47]. Therefore, the increased propensity for hyperoxia-induced increased MPTP opening and decreased ΔΨm in AT2 from mice with C57 mtDNA compared to mice with C3H mtDNA noted in this study indicates that differences in AT2 apoptosis modulated by differences in mtDNA may be a contributory mechanism for decreased alveolarization and increased lung injury in these mice. Interestingly, Cyclosporin A, which is an inhibitor of MPTP opening, was noted to protect against hyperoxic lung injury in adult but not neonatal mice [48,49]. Since glutamate-induced NMDA receptor dependent calcium influx is known to increase MPTP opening, the differences in downregulation of glutamatergic signaling that we noted in the lung transcriptome of only certain mito-nuclear combinations suggests that inhibition of MPTP opening may be more effective for individuals with certain mtDNA haplogroups but not others [50].

Since the lung functions in an oxygen-rich environment it is continuously exposed to oxidative stress. Excessive ROS that overwhelms mitochondrial anti-oxidant defenses has been shown to increase mitochondrial membrane permeability and lead to cell death [46,51]. While we did not specifically analyze ROS generation from AT2 cells in this study, we have previously shown that NMLF O_2_^−^ generation is higher in mice with C57 mtDNA compared to mice with C3H mtDNA [16]. In this context, the increased H_2_O_2_ and protein carbonyl content found in the lungs of all mouse strains exposed to hyperoxia in this study indicates that AT2 cells were exposed to an environment in which oxidant stress was increased. Additionally, the higher content of oxidative stress markers such as O_2_^−^ and MDA in the lungs of mice with C57 mtDNA also suggests that AT2 and NMLF from these mice were exposed to higher levels of oxidative stress compared to mice with C3H mtDNA. Furthermore, in addition to the bioenergetic and metabolic state of the cell, MPTP opening is also influenced by mitochondrial ROS. Oxidant damage also markedly reduces activity of enzymes such as SIRT3 and aconitase (both of which were decreased in hyperoxia-exposed mice with C57 mtDNA compared to mice with C3H mtDNA) which play critical roles in the maintenance of mtDNA integrity and SIRT3 deficiency in AT2 has been shown to promote lung fibrosis through increased mtDNA damage [52,53].

In addition to their well-recognized role in bioenergetics, metabolism and redox biology, mitochondria also mediate - and are targets of - inflammatory responses to a variety of noxious stimuli. This is because they are evolutionary remnants of ancestral bacteria and contain several components such as mtDNA that can act as potent damage associated molecular patterns (DAMPs) to recruit inflammatory/immune responses [54]. Mitochondrial role in these processes has been implicated in lung diseases such as COPD [55]. One specific mechanism of mtDNA-DAMPs mediated inflammation involves signaling through stimulator of interferon response cGAMP interactor 1 (STING1) which consequently increases synthesis of cytokines such as IFN-β, IL-6 and TNF-α, all of which were found to be higher in mice with C57 mtDNA compared to mice with C3H mtDNA, implying that damaged mtDNA extruded from AT2 cells may contribute to the increased inflammatory cytokine response to hyperoxia noted in the C57 mtDNA group. Finally, we noted that cytokines related to eosinophil activation varied with mtDNA carried by mice. Previously, we have shown that eotaxin may be a marker for BPD risk, while others have found that asthmatic individuals with certain mitochondrial haplogroups have increased IgE levels but not others [5,56]. Taken together, such evidence suggests that mtDNA variations may modulate BPD risk through differences in eosinophilic inflammation.

Mitophagy, which is a mitochondrial quality control mechanism, degrades populations of low-potential mitochondria to restore cellular homeostasis and ensure continued survival. Therefore, increases in depolarized mitochondria, as seen in AT2 from mice with C57 mtDNA compared to mice with C3H mtDNA is indirect evidence that mitophagy may have failed or become insufficient in mice with C57 mtDNA. Impaired PINK1/Parkin-mediated mitophagy contributes to the accumulation of damaged mitochondria and has been implicated in disorders such as COPD and IPF [57,58]. Additionally, activation of mitophagy through pharmacological agents such as rapamycin and urolithin A has been demonstrated to reduce oxidative stress, attenuate inflammation, and promote lung epithelial cell survival. Previous studies have found that both inherited and acquired mtDNA mutations are associated with differential expression and regulation of mitophagy-associated proteins such as PINK1 [59,60]. Therefore, our findings that mice with C57 mtDNA have decreased mitophagy in AT2 and NMLF suggests that such variable modulation of lung mitophagy based on mtDNA haplogroup may be a contributory factor to the differences in BPD risk noted in infants from diverse racial groups and requires further investigation.

Polymorphisms that differentiate C57BL6 and C3H/HeN mtDNA have been identified in genes that encode for the ND3 subunit of complex I, subunit III of complex IV and tRNA^Arg^ [16]. While no consistent differences were noted in C-I OCR, the lower C-IV OCR that we noted between hyperoxia-exposed permeabilized AT2 from mice with C57 mtDNA compared to mice with C3H mtDNA indicates that mtDNA variants may contribute to differences in lung injury through modulation of AT2 bioenergetic function. Mitochondrial-nuclear interactions have also been noted to occur via “retrograde signaling” through which changes such as decreased ΔΨm, ATP, mt-ROS and other mitochondrial signals modify nuclear responses to external and internal stimuli [61]. Our observations revealed striking similarities in the lung transcriptome changes between WT mice strains and their corresponding MNX mice, suggesting that the response to hyperoxia in the lungs of newborn mice is predominantly regulated by nuclear genes, which is to be expected. However, the differences we noted such as higher degree of upregulation of inflammatory pathways as well as downregulation of glutamatergic signaling in mice with one set of mito-nuclear combination but not the other implies that similar mito-nuclear interactions may also be operative in some infants in whom mitochondrial retrograde signaling pathways may have more profound and/or variable effects on nuclear-mitochondrial interactions than other infants, leading to differences in their predisposition to lung injury.

Finally, genetic variation related to lung development, and immune responses are known to contribute to individual and racial/ethnic differences in respiratory outcomes in high-risk premature infants [62]. Since mtDNA haplogroups segregate largely by ethnicity, it is possible that mtDNA haplogroups could influence BPD risk and pathogenesis. Endothelial cell bioenergetics varies with mtDNA haplogroups and we have previously shown that mitochondrial bioenergetic function is decreased in HUVEC from both term and prematurely born Black infants (who most commonly carry the L mitochondrial haplogroup) compared to White infants (who most commonly carry the H mitochondrial haplogroup) [14,63]. Our findings that infants with different haplogroups had variable levels of SIRT3 is supported by similar findings of differences in SIRT3 expression in cybrids with different mtDNA haplogroups exposed to oxidative stress in a previous study. The observation that Black infants with BPD had evidence of decreased mitophagy in HUVEC at birth as well as in TA samples obtained at 28 days of life when compared to White infants who also had BPD suggests that Black infants may have a predisposition for mitochondrial dysfunction and potentially benefit more from mitochondrially-directed therapies compared to White infants [64].

Our study has a few limitations. While we measured mitochondrial bioenergetic function, we did not test glycolytic capacity in cells which may also modulate MPTP opening through pH dependent effects as has been previously noted [46]. We were also unable to perform immune-TEM of lung sections which could have help quantify mitophagy more accurately. However, we believe that the multiple converging lines from evidence including transcriptomic changes that were confirmed using wet-lab qPCR as well as direct examination of mitophagy through flow-cytometry based methods in lung cells makes up for this limitation. Similarly, though we believe that the multiple analytes we used to measure ROS content in lungs serve as convincing evidence, fluorescence-based methods to detect ROS carry their own limitations, which we hope to address through more robust analysis of ROS generation in specific lung cell populations using techniques such as electron paramagnetic resonance. Finally, we could only explore pathways related to inflammation and mitophagy but not others such as glutamatergic signaling or the *microRNAs related to cancer* pathway which were upregulated only in mice with C3HWT mtDNA in this study. We hope to address these limitations and also obtain information regarding the specific mtDNA haplogroups (instead of using self-reported race) of the human cohort in subsequent studies.

Despite such limitations, to our knowledge, this is the first study that establishes mechanistic links between mtDNA variations and risk for neonatal lung injury. Our results lend support to a framework for neonatal lung injury in which differences in mtDNA, which are functionally inconsequential during basal states, lead to differential levels of mitochondrial bioenergetic function, ROS generation and mitochondrial membrane integrity that subsequently leads to variable degrees of mitochondrial quality control and inflammatory lung injury and lung cell function and finally culminates in causing varying risk for dysregulated lung development and BPD in infants with different inherited mtDNA haplogroups.

In conclusion, several determinants of health and disease are known to be modified by mitochondrial-nuclear interactions and a deeper understanding of the magnitude and mechanisms underlying such interactions could help improve our understanding of the basis of complex diseases with multifactorial etiology such as BPD. Future studies should be aimed at investigating mechanisms such as epigenetic regulation of nuclear gene expression through retrograde signaling by mitochondria [65]. While such investigations are likely to be complicated by the high degree of heteroplasmy that is noted in mtDNA at the individual, tissue, and cellular levels, understanding the underlying mechanisms of mitophagy dysregulation in specific lung cell populations in mice and accessible cells such as MSCs, HUVEC and platelets from infants can pave the way for the development of novel therapeutic strategies aimed at restoring mitochondrial homeostasis and improving lung health.

## 4. Methods

### 4.1. MNX mice BPD model and sample collection from mice and ELBW infants

C57 and C3H WT and MNX strains were obtained from colonies maintained at the University of Alabama at Birmingham (UAB) and validation of mitochondrial haplotypes was done using single nucleotide polymorphisms as described previously [66]. Newborn mice were exposed to ambient oxygen (Room Air or RA; 21% O_2_) or hyperoxia (85% O_2_) from postnatal days 4-14 (P4-14) to create lung injury following a well-established protocol that has been previously described [67,68]. Lungs obtained from mice following RA or hyperoxia exposure were homogenized to obtain tissue lysate and isolated mitochondria or used to isolate NMLF and AT2 as described previously [69].

HUVEC were isolated following a previously described protocol from ELBW infants at their birth [14,70]. TA samples were collected through endotracheal tubes of ventilated ELBW infants by instilling 0.5 mL of 0.9% saline followed by suctioning of residual fluid with a 5F catheter as part of routine clinical care protocols followed for these infants in our NICU and centrifuged at 400 rpm for 5 min to isolate cells. Both HUVEC and cell pellets isolated from TA were cryopreserved in liquid nitrogen until later use.

### 4.2. AT2 bioenergetic function

AT2 mitochondrial respiration assays were performed using high resolution respirometry by measuring oxygen consumption in an Oxygraph-2k two-channel respirometer (Oroboros). 70% ethanol was run in both chambers for a minimum of 30 minutes and the chambers were calibrated after a stable air-saturated signal was obtained before every experiment. Reactions were conducted at 37°C in a 2 ml chamber containing air-saturated mitochondrial respiration medium (Oroboros, 60101-01) with 1×10^6^ cells/mL under continuous stirring. Intact cells were used to measure basal respiration as well as maximal (uncoupled) rates measured after addition of 0.5 uM FCCP. Cells permeabilized with digitonin (optimized using a titration assay and cytochrome c) were used to measure state 3/complex I respiration after addition of 5 mM malate/15 mM glutamate followed by inhibition of complex I with 0.5 μM rotenone, state 3/complex II respiration after addition of 2.5mM ADP/10 mM succinate/ 200 μM palmitate conjugated with 0.25% BSA followed by inhibition of complex III with 5um Antimycin A, complex IV respiration after addition of 2mm ascorbate/0.5 mM N,N,N’,N’-tetramethyl-p-phenylenediaminepalmitylcarnitine (TMPD) followed by inhibition with 5mM sodium azide and finally, state 4o respiration after addition of 1.3 μM oligomycin. RCR was calculated as a ratio of state3/state4o respiration [71]. A colorimetric ATP assay kit (Abcam, ab83355) was used to measure ATP content in lung tissue using a SpectraMax^®^ i3x reader according to manufacturer’s instructions. A minimum of 12 mice per group of either sex were used for the AT2 cell isolation for these experiments.

### 4.3. MPTP opening

MPTP opening in AT2 was measured using ionomycin-induced calcein signal quenching as described previously. Briefly, AT2 grown in 8-well chambered coverslips (5 x 10^4^ cells/well) and exposed to RA or hyperoxia for 48 hours were washed with modified Krebs Ringer Buffer (KRB) containing 135 mM NaCl, 5 mM KCl, 0.4 mM KH_2_PO_4_, 1 mM MgSO_4_·7H_2_O, 20 mM HEPES, and 5.5 mM glucose and 1 mM CaCl_2_ at pH 7.4. Next cells were incubated with 1mM Calcein (Abcam, 148504-34-1) for 15 min at 37°C in 5% CO_2_ and washed with modified KRB to remove excess dye. Basal images were first obtained using a Nikon A1R confocal microscope with a FITC excitation filter followed by further imaging after the addition of 10 µM ionomycin to stimulate MPTP opening. Rate of Calcein signal quenching after the first minute of ionomycin in a minimum of 5 fields from cells obtained from a minimum of 6 mice per group was used to measure and compare MPTP between the different groups as described previously [72–74].

### 4.4. Mitochondrial membrane potential

Mitochondrial membrane potential (ΔΨm) of AT2 was measured using TMRM (Thermofisher, 134361) according to manufacturer’s instructions. Briefly, AT2 were seeded chambered coverslips (5 x 10^4^ cells/well) were exposed to RA or hyperoxia for 48 hours were incubated with TMRM in the dark for 30 min at 37 °C and washed with PBS to remove excess dye. Fluorescence images were captured using a TE2000U microscope (Nikon) fitted with a TRITC filter and a QiCam Fast Cooled high-resolution CCD camera and mean fluorescence intensity of TMRM of 10 images per mice from a minimum of 12 mice per group was analyzed using Fiji v2.9 [75].

### 4.5. MtDNA damage analysis

To quantify mtDNA damage genomic DNA was extracted from AT2 using the QIAamp DNA mini kit (Qiagen, 51304) and quantified using the PicoGreen assay kit (Invitrogen, P7589). A 16059-bp region of the NADH5/6 genes in the mouse mtDNA genome was amplified via qPCR by using primer set M13597 FOR (13597 to 13620 bp) and 13361 REV (13361 to 13337 bp. Copy number differences were normalized by amplifying a shorter 80 bp segment of the mtDNA using the primers 13597F (5′CCCAGCTACTACCATCTTCAAGT) and 13713R (GATGGTTTGGGAGATTGGTTGAT GT3. The resulting gels were dried, visualized and densitometry was performed using Fiji v2.9 to normalize the 16059-bp QPCR product with the 80-bp qPCR product and DNA lesion frequency/10 kb of DNA was quantified as described previously using a minimum of 12 mice per group [76].

### 4.6. Aconitase activity

Mitochondrial aconitase activity was measured using an aconitase activity kit (Abcam, ab83459) in mitochondria isolated from AT2 from mice suspended in the provided assay buffer provided and then incubated for 1 h with the isocitrate substrate mixture at 25°C. Absorbance at 240 nm was recorded for 30 min using a SpectraMax^®^ i3x reader. Catalytic activity of aconitase was determined by measuring the rate of formation of cis-aconitate as detected by increase in the absorbance per manufacturer’s instructions from a minimum of 12 mice per group.

### 4.7. Oxidative stress markers

O_2_^−^ content was measured by incubating freshly isolated lung homogenates with 5 μM/L lucigenin (Millipore Sigma, M8010) and 500 μM/L of NADH or NADPH for 10 min at 37 °C followed by chemiluminescence measurements using a SpectraMax^®^ i3x plate reader at 1 min intervals over a 5 min period [77]. H_2_O_2_ content was measured by incubating lung homogenates with 100 μM/L of Amplex Red and horseradish peroxidase (Invitrogen, A22188) for 30 min at 37 °C, followed by measurement of absorbance at 560 nm in the supernatants using a SpectraMax^®^ i3x plate reader [14]. Lung MDA content was measured by centrifugation of freshly homogenized mice lung tissue and incubating the supernatants with thiobarbituric acid (Abcam, ab118970) at 95°C for 1 hour followed by cooling to RT and measuring absorbance on a SpectraMax^®^ i3x plate reader at 532 nm according to manufacturer instructions. To measure protein peroxidation, protein samples extracted from lung homogenates were incubated with 2,4-dinitrophenylhydrazine (Abcam, ab126287) followed by precipitation and resolubilization with trichloroacetic acid, acetone and guanidine and absorbance measurements at 375 nm using a SpectraMax^®^ i3x plate reader according to manufacturer instructions.

### 4.8. Cytokine content analysis

Cytokine protein concentration was measured in lung homogenates that were incubated in V-Plex Proinflammatory Panel 1 Mouse Kit (Mesoscale Discovery, K15048D) multi-well plates pre-coated with antibodies for IL-1β, IL-6, TNF-α, IL-2, IL-12p70, IFN-γ, KC/GRO (CXCL1), IL-4, IL-5, and IL-10 and allowed to react for 2 hours according to the manufacturer’s instructions. After detection antibody was added to each well and allowed to react for 1 hour, plates were analyzed on an MSD QuickPlex SQ 120 instrument (MSD, AI0AA-0) and Discovery Workbench v4.0 was used to calculate cytokine concentrations using a linear regression analysis of the standard curve as described previously from a minimum of 12 mice per group [78].

### 4.9. Lung transcriptome analysis

Total RNA samples extracted using RNA STAT-60 (AMSbio, CS-111-200) from mice lung homogenates were sent to Genewiz (Azenta Life Science) for sequencing and library preparation using polyA selection and the Illumina Hi-Seq platform to generate paired-end unstranded fastq files. Raw read quality was evaluated using FastQC v0.11. Adapter and low-quality bases below a quality score of 30 were trimmed from raw RNA sequencing reads using BBDuk2 v38.73 and the trimmed reads were mapped to the Mus musculus GRCm39 reference genome available on ENSEMBL using the STAR aligner v.2.5.2b to generate raw counts files. Batch correction was performed using CombatSeq v3.3.6 and, after normalization using variance-stabilizing transformation (VST), differentially expressed genes (DEGs) were identified using DESeq2 using LFC > 2 and FDR p-value < 0.05 [79]. The online platform IDEP (v1.0) and the R package ClusterProfiler was used to perform pathway analysis of all genes with LFC values > 1 using Generally Applicable Gene-set Enrichment (GAGE) and the Kyoto Encyclopedia of Genes and Genomes (KEGG) database to identify pathways that were differentially regulated with LFC > 2 and FDR p-value < 0.05 [73,80]. RNA from 3 mice per treatment group were used to perform these analyses.

### 4.10. TEM imaging

Fresh mouse lungs were minced into fine pieces and fixed using 2.5% glutaraldehyde in PBS at 4 °C. Tissues were post-fixed in 1% osmium tetroxide in 1% K_4_Fe(CH)_6_, dehydrated through graded concentrations of graded ethyl alcohols from 50 to 100%, embedded using EMbed 812 (Electron Microscopy Sciences, EMS #14120), sectioned with a diamond knife, placed on copper grids, stained with uranyl acetate/lead citrate and visualized with a Tecnai Spirit 120kv TEM (Nanoimaging Services). Digital images were taken with an AMT BioSprint 29. A minimum of 20 images from at least 6 mice per group were used to quantify autophagic vacuoles and mitochondrial area per cell using Fiji v2.9.

### 4.11. Western blots

30 to 60 µg of protein obtained from AT2 were separated on 4-20% Criterion TGX Gels (Bio-Rad. Cat. No. 5678093), transferred to PVDF membranes (Trans-Blot Turbo System, Bio-Rad) and probed with antibodies against PINK1 (Invitrogen, PA1-16604), Parkin (Invitrogen, 702785), ETC complexes I-V (total OXPHOS cocktail, Abcam ab110413) and SIRT3 (Invitrogen, PA5-86035) with β-Actin (Cell Signaling Technology, 5125S) as loading control. A similar procedure was used to probe protein samples obtained from HUVEC from ELBW infants with antibodies against PINK1, Parkin and SIRT3 with β-actin as loading control. All membranes were visualized with ChemiDoc imaging System (Bio-Rad) and densitometry of band intensities was performed using Fiji v2.9. All WBs were carried out using 3 independent samples per group.

### 4.12. QPCR for RNA

RNA was isolated from AT2, HUVEC and TA-derived cell pellets using RNeasy Plus Mini Kits (Qiagen, 74136). cDNA reverse-transcribed from 1 µg of this RNA using iScript Reverse Transcription Superrmix (Bio-Rad, 1708840) was used as a template for performing qPCR using TaqMan Fast Advanced Master Mix (Thermo Fisher Scientific, 4444556) with TaqMan gene expression assays for PINK1 (mice - Mm01348629_m1, human - Hs00260868_m1), Parkin (mice - Mm01323528_m1, human - Hs01038322_m1), SIRT3 (mice - Mm00452131_m1, human - Hs00953477_m1) and PGC-1α (mice - Mm01208835_m1) according to the manufacturer’s instructions on the Bio-Rad CFX96 system with normalization performed using eukaryotic 18S rRNA Endogenous Control (Thermo Fisher Scientific, 4310893E). qPCR data was analyzed by applying the comparative Ct method (ΔΔCT). The results were presented as the fold change in mRNA expression for targeted genes relative to controls. All qPCRs were carried out using 3 independent samples per group.

### 4.13. Flow cytometry

Mitophagy in freshly isolated AT2 cells was detected using the Mitophagy detection kit® which contains a Mtphagy and Lysosomal dye (Dojindo Molecular Technologies, MD01-10). 10,000 AT2 cells per sample were treated with 100 nM Mtphagy Dye for 15 min. After washing away excess with PBS, the cells were exposed to RA or hyperoxia for 12 hours, following which they were treated with 1 µM Lysosomal Dye and incubated for 10 min. After washing excess with PBS, cells were imaged (Ex/Em – 561 and 650 nm for MtPhagy dye and 488/550 nm for Lysosomal dye) using an Amnis ImagestreamX MKII imaging flow cytometer (EMD Millipore). Appropriate unstained cell and single-color cell controls were used to generate a compensation matrix which was used to convert raw image files to compensated image files and the co-localization wizard from IDEAS® v6.0 (Amnis) was used to calculate BDS scores for Mtphagy and Lysosomal dye in 10,000 cells/sample from a minimum of 3 mice per group.

To measure mitophagy in HUVEC were seeded in 6-well plates (8 x 10^5^ cells/well) and transfected with 5ug/well of pCHAC-mt-mkeima plasmids (Addgene plasmids #72342) at a ratio of 1:1.5 plasmids: lipofectamine 3000 (Invitrogen, L3000001) overnight. Next, these were then treated with RA or hyperoxia for 12 hours, removed from wells using trypsinization, washed with PBS and resuspended in 100 µl of 1:1000 near-IR live/dead stain (Invitrogen, L10119) for 10 minutes. Following this, cells were washed again with PBS and suspended in 1 mL of 10% PBS for flow cytometry. An LSR Fortessa flow cytometer with Violet (405 nm) and Yellow/Green (561 nm) was used to measure cytosolic (mitochondrial) and lysosomal mt-Keima fluorescence. Gating was carried out to separate single live cells from debris and dead cells using untransfected and unstained cells as controls, following which the ratio of cytosolic to lysosomal fluorescence was calculated for 20000 events/sample from a minimum of 3 mice per group as described previously [81].

### 4.14. Statistical analysis

Data are presented as median ± interquartile range (IQR). Normality was tested using the Shapiro-Wilk method. Differences between groups were analyzed by one-way or two-way ANOVA according to experimental design followed by Tukey’s post hoc test for normally distributed data, and Kruskal Wallis test for non-normal data. Student-t test or Mann-Whitney U test were used for individual comparisons between groups as appropriate. A two-tailed p-value of less than 0.05 (95% confidence level) was considered statistically significant and Bonferroni method was used to adjust for multiple comparisons. All statistical analyses were performed using R v4.2.7 [82].

### 4.15. Study Approvals

All protocols were approved by the Institutional Animal Care and Use Committee and the Institutional Review Board of the University of Alabama at Birmingham (UAB) and consistent with the Public Health Service Policy on Human Care and Use of Laboratory Animals (Office of Laboratory Animal Welfare, 2002).

## Author contributions

JK, SWB and NA conceived and designed the study; RL, JK, TJ, BMV, and SH performed experiments; JK, RL, SWB and NA analyzed the data; JK, NA, MNG and SWB interpreted the results of experiments; JK prepared the figures; JK and NA drafted the manuscript; JK, RL, BMV, SH, TJ, MNG, SWB and NA revised and approved the final version of manuscript.

